# Pangenome-conserved k-mers enable accurate de novo and somatic tandem repeat mutation calling from long reads

**DOI:** 10.64898/2026.11.23.762899

**Authors:** Ellie Rahm Kim, Heng Li

## Abstract

Although long reads span most tandem repeats, calling de novo or somatic tandem repeats remains challenging due to sequencing and alignment errors. Here, we developed TRumut, which uses a pair of pangenome-conserved k-mers around each tandem repeat to measure length variations across reads and samples. TRumut does not need read alignment and thus avoids alignment artifacts. We further characterized recurrent sequencing errors in tandem repeats and established thresholds to separate mutations from artifacts. TRumut successfully resolved de novo and somatic mutations across a four-generation pedigree and ten cancer cell lines, accurately classifying events by clonality and identifying recurrent loci.

## Background

Tandem repeats are genomic regions in which a sequence motif recurs in succession. Repeats with 1-6 bp motifs are called short tandem repeats (STRs), and those with longer motifs are called variable number tandem repeats (VNTRs). A repeat allele is characterized by its number of repeat units and its internal sequence [1,2], and it can change in either dimension. In the most mutable STRs, these changes occur at rates of 10^-4^ to 10^-3^ per locus per generation [3], four to five orders of magnitude higher than the ∼10^-8^ per-site rate of SNVs [4]. Repeat alleles therefore differ between individuals, between generations, and between the cells of a single individual. Changes present in every cell of an individual but in neither parent are termed de novo mutations, and those confined to a subset of an individual’s cells are termed somatic mutations.

Tandem repeat mutations may be causal forces in human diseases. In Huntington’s disease, individual neurons undergo CAG repeat expansions in the *HTT* gene and die only after their own repeat passes approximately 150 units [5]. Comparable somatic instability is documented in myotonic dystrophy [6], fragile X syndrome [7], *C9orf72* ALS/FTD [8], and the spinocerebellar ataxias [9]. In cancer, 160 repeat expansions across 2,622 tumor genomes showed signs of selection, recurring within specific cancer types and clustering at regulatory elements [10]. One of these expansions was present in a third of renal cell carcinomas, and targeting it with a small molecule reduced cancer cell proliferation [10]. Even in healthy individuals, repeats constantly mutate somatically [11], and repeat variation alters the transcriptional output [12]. In many of these settings, only a single cell or a small population of cells holds the consequential allele. Understanding the role of tandem repeats in disease therefore requires resolving somatic repeat alleles.

Short-read sequencing does not provide this resolution. A read must span the repeat and enough flanking sequence to tell which locus it came from. Short reads can therefore only provide accuracy in short repeat tracts. For longer repeats, repeat lengths are instead estimated by aggregating partial reads, which blurs the genotype and erases any allele carried by a minority of cells [10,13,14]. Thus, most pathogenic expansions and VNTRs cannot be studied with short reads [15,16]. In theory, long reads lift that constraint. They routinely span entire repeat loci [1], so the allele sequence can be read off each read rather than estimated. A read also comes from a single DNA molecule and therefore a single cell, which is the resolution at which mutations arise.

In practice, telling apart true somatic tandem repeat mutations from errors has been a major limiting factor. First, tandem repeat callers rely on alignment to establish where each repeat begins and ends [1,17–21]. However, alignment is often unreliable in tandem repeat regions where boundaries may be misplaced, yielding incorrect alleles. Second, although HiFi reads are 99.8% accurate, 92% of their residual errors are homopolymer indels [22], the simplest form of tandem repeats. Consistent with this, germline benchmarks find HiFi worse than short reads in homopolymers and hint it is no better at other repeats [23,24]. However, profiling this error across tandem repeats requires accurate repeat calls that alignment cannot deliver. Its extent thus remains unknown.

Existing callers, such as TRGT [1], Straglr [17], and LongTR [18], manage these pitfalls by collapsing spanning reads into one or two consensus alleles per locus. This works for germline genotyping but discards the read-level information for somatic alleles. Meanwhile, methods built specifically for somatic repeat variation report summary statistics of instability rather than the mutations themselves [25–27]. Two methods call individual somatic events, but with limited scope: MSMuTect detects somatic microsatellite indels in tumor-normal exomes [28], and prancSTR infers a single high-frequency somatic allele per locus without a matched control [11]. Both infer the mutant allele statistically from short reads rather than reading it from a molecule, and neither extends beyond short microsatellites. Therefore, little is known about tandem repeat mutations at the level of individual molecules and events.

To address this gap, we developed TRumut, an alignment-free caller that “reads off” the allele sequences from long reads (Fig. 1a). TRumut sidesteps alignment by anchoring each repeat to ultraconserved single-copy k-mers we cataloged across the human pangenome. It does not collapse reads into a consensus, so every allele in a sample remains detectable, including those carried by a small fraction of cells.

**Figure 1.**
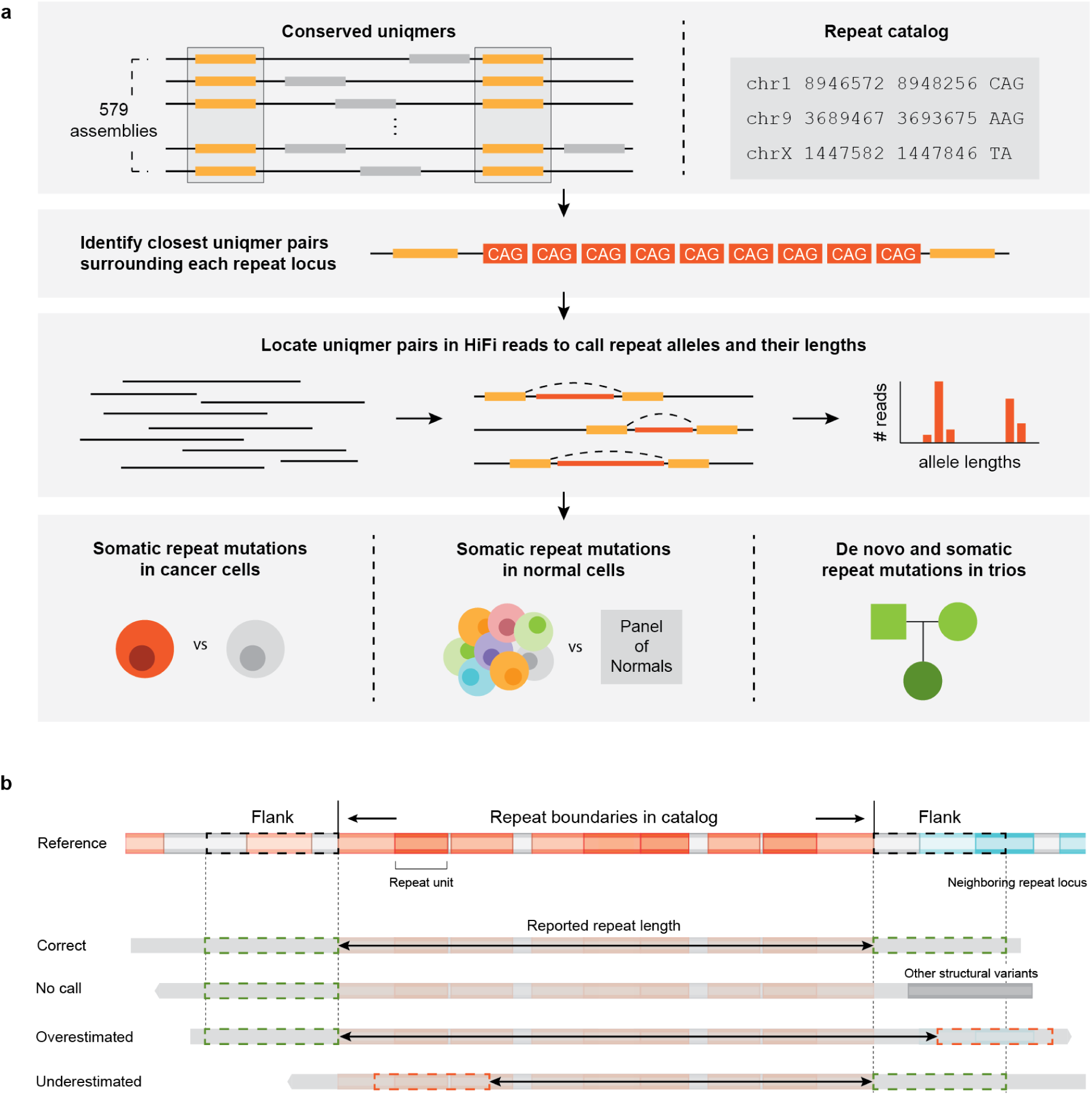
Alignment-free calling of tandem repeat alleles with TRumut. a. Overview of the TRumut workflow. Top: uniqmers (yellow boxes) occur exactly once per haplotype and are conserved across 579 haplotype assemblies (left). Each catalog locus (right) is annotated with the closest flanking uniqmer pair (middle; here a CAG repeat). Middle: in each HiFi read (black lines), TRumut locates both uniqmers by exact match and takes the intervening sequence (red) and its length as that read’s allele. Bottom: per-read alleles are compared against a matched normal (cancer), a panel of normals (unmatched samples), or the parents (trios) to call mutations. b. Failure modes in alignment-based repeat calling. Alignment-based callers locate a repeat by aligning its two flanks and report the sequence between them, so each call is only as reliable as the two flank placements. Top: reference sequence at a repeat locus, with the catalog interval marked above and the flanks boxed. Red, repeat units, lighter and darker for units differing in sequence; gray, non-repetitive sequence; cyan, a neighboring repeat. Below: reads shaded by the reference sequence they carry. Each flank placement is boxed, green if correct and red if not, and the reported length is the arrow between them. Correct: both flanks placed correctly. No call: other structural variants in a flank prevent its alignment, so nothing is reported. Overestimated: a flank aligns to a similar sequence in the neighboring repeat. Underestimated: the read does not reach the left flank, but the flank is aligned on flank-like sequence inside the repeat instead of no call being reported.

Eliminating alignment errors lets us measure long-read sequencing errors in tandem repeats directly. By calling the same genome against itself across replicate runs, chemistries, and platforms, we obtained, to our knowledge, the first genome-wide, read-level profile of long-read error in tandem repeats. We set calling thresholds from these distributions. In a four-generation pedigree, we recovered de novo and postzygotic repeat mutations and followed individual alleles that mutated repeatedly as they were inherited. Across six tumor-normal pairs, we found somatic mutations that differed in clonality between lines, recurred at shared loci, and reached long VNTRs, a class no existing somatic caller reports. Calling all 10 cell lines against a population panel showed microsatellite instability destabilizing STRs but not VNTRs.

## Results

### Alignment-free tandem repeat allele calling with TRumut

Repeat calling requires two decisions for each read: which locus the read derives from, and where the repeat lies within it. Standard tools make both by alignment. They take the repeat’s boundaries from a catalog annotated on the reference genome, then locate those boundaries in each read by aligning the reference sequence on either side of them. However, this is circular: the caller needs the repeat’s boundaries to measure its length, and it finds those boundaries by aligning to sequences that the repeat itself has made ambiguous. Reads of one haplotype end up with different boundaries and lengths (Fig. 1b; Additional file 1: Figs. S1 and S2). Since a mutation is often called from a length difference, these shifts are indistinguishable from real mutations.

We instead leverage “uniqmers,” which we define as 31-mers that occur exactly once per haplotype and are conserved across the human pangenome (Fig. 1a; Methods) [29]. Since a uniqmer marks a single, rarely varying position, a read carrying one can be placed in the genome unambiguously. Likewise, a pair of uniqmers flanking a repeat tells us where the repeat lies within the read. We call each flanking uniqmer an anchor and the two together an anchor pair. An important advantage of these anchors is that they are consistent across individuals unlike catalog boundaries defined on the reference genome only: a boundary drawn on GRCh38 may have no counterpart in a genome with a different repeat tract, but a uniqmer, defined on the pangenome instead, marks the same position across genomes.

We first validated what fraction of repeats uniqmers could anchor. We found that 34% of all 31-mers in GRCh38 occur exactly once in at least 98% of 579 haplotype assemblies (Additional file 1: Fig. S3a) [29]. At that density, the nearest uniqmer lies a median of 26 bp from each repeat boundary (Additional file 1: Fig. S3b). We could not anchor only 0.3% of repeats, nearly all near telomeres and centromeres. Our final catalog comprised 2,254,054 anchored regions covering 3,433,388 tandem repeats; one region may contain two or more repeats if they are close and there are not uniqmers between them.

For every read spanning an anchored region, we record the sequence between the anchors, its length, and the number of k-mers matching the repeat motif (Fig. 1a). It was possible that the interval included some non-repetitive sequence lying between the repeat and the anchors. To avoid mistaking an insertion or deletion in these regions as a repeat mutation, we also recorded the count of motif k-mers for each read (Methods). A true repeat mutation shifts length and motif content together, while a flanking indel shifts length alone.

Throughout, we use the term “allele” to denote a set of reads at a locus whose inter-anchor lengths agree within a few base pairs, annotated by its most frequent length. A “candidate” is an allele in a test sample that differs from every allele in a base sample which could be the parents in a trio, the normal sample in a tumor-normal pair, or a technical replicate for measuring sequencing error rates. An “event” is a candidate that passes the thresholds derived below. Finally, a “lineage” is an allele tracked across generations through pedigree descent.

### HiFi technical replicates disagree at a tenth of tandem repeat loci

With alignment removed, we expected sequencing to be the dominant source of error. To measure this error, we turned to technical replicates of the Genome in a Bottle sample HG002 (datasets and coverage in Fig. 2). Since technical artifacts are typically randomly distributed across runs, alleles unique to a replicate provide an approximation of platform error. We counted such discordant alleles three ways. First, we included all alleles, including those supported by single reads and differing by ≥3 bp in length. Second, we restricted counts to alleles supported by at least two reads of identical length, which excludes errors confined to one molecule. Third, we evaluated alleles differing by 7 bp or longer from any allele in the comparison dataset.

**Figure 2.**
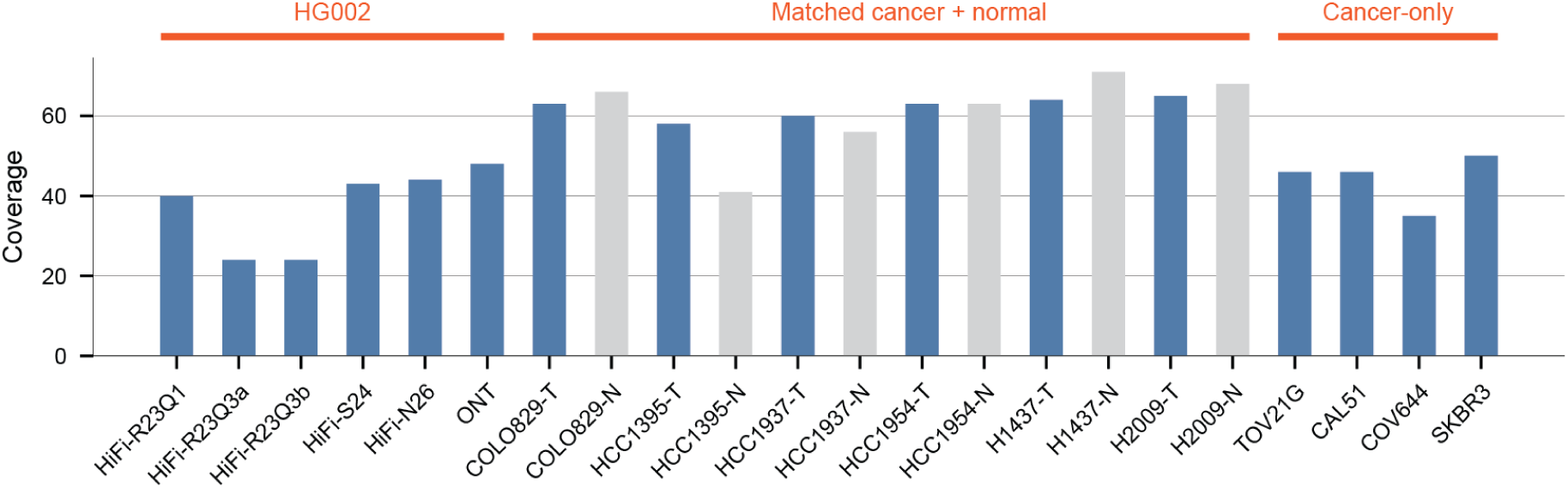
Sequencing coverage of the datasets analyzed in this study. Mean read depth of each dataset before downsampling, grouped as HG002 technical replicates, cancer cell lines with a matched normal, and cancer cell lines without. Matched normals (-N) in gray, all others in blue. Download locations and accessions are given in Availability of data and materials.

We first compared two SMRT runs of a single library, sequenced a week apart on the same instrument (HiFi-R23Q3a, HiFi-R23Q3b), together with an earlier and independent run (HiFi-R23Q1). We downsampled all three to equal coverage (24x) and compared them at 1,810,012 loci covered in all three (Methods). Across all three datasets, 228,785 loci, 13% of those analyzed, were discordant in at least one comparison (Fig. 3a). Requiring at least two reads of the same length left 23,864 loci (1%), and requiring an offset of more than 6 bp left 72,891 (4%) (Additional file 1: Fig. S4a, b). Both counts are far more than what one would expect from two runs of one genome. Even the two runs of one library disagreed: HiFi-R23Q3a carried 101,351 alleles absent from HiFi-R23Q3b, and HiFi-R23Q3b carried 86,276 absent from HiFi-R23Q3a.

**Figure 3.**
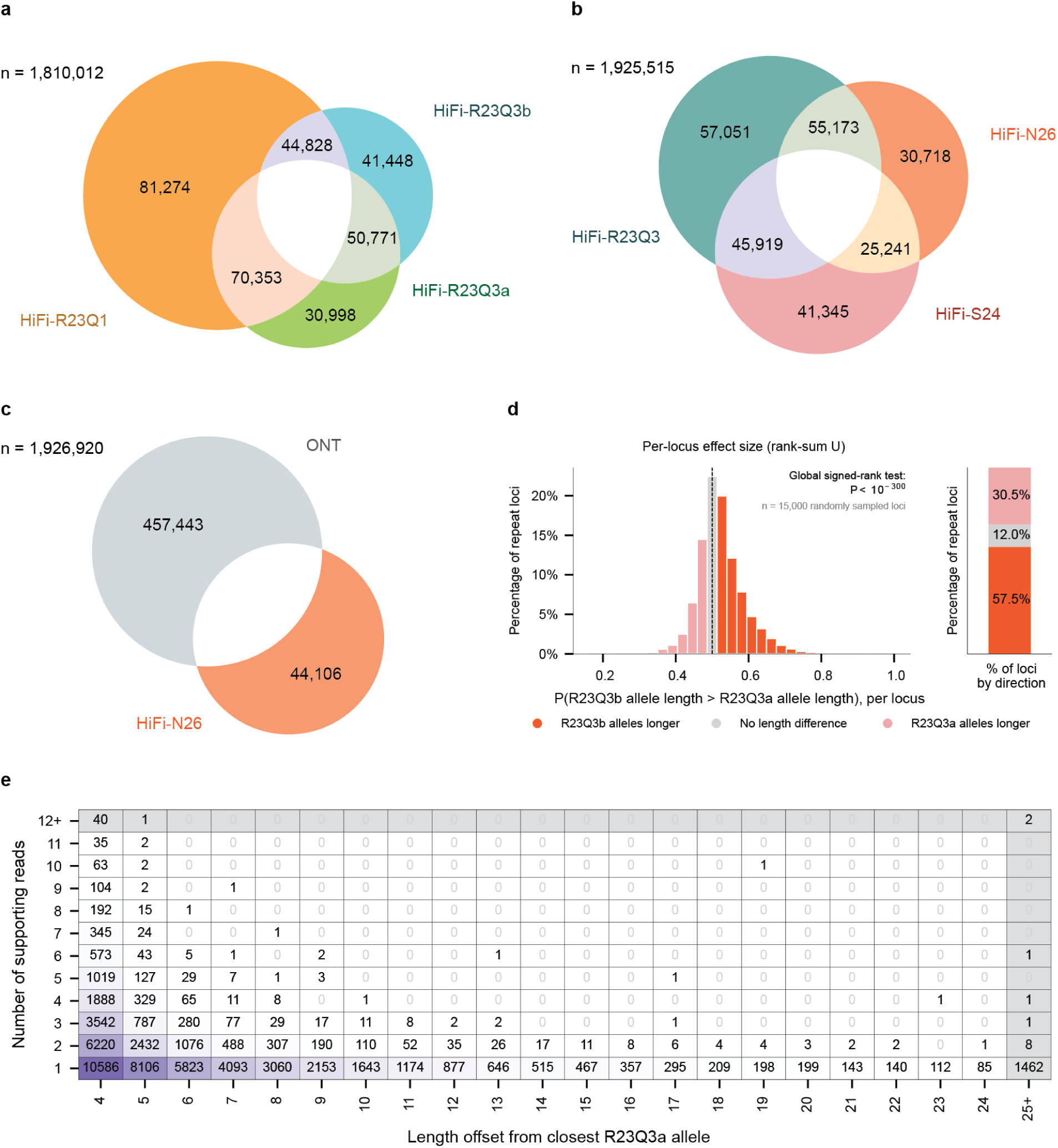
Error profile of HiFi sequencing in tandem repeats, measured from technical replicates. a-c, Loci with an allele detected in one dataset but absent from another, identified with the mutual module (Methods). n, loci covered by at least 10 reads in every dataset. Homopolymers were excluded, and datasets were downsampled to equal coverage. A locus appears in more than one segment when it carries alleles unique to more than one dataset. a. An earlier run (HiFi-R23Q1) and two runs from the same library (HiFi-R23Q3a, HiFi-R23Q3b), at 24x. b. HiFi-R23Q3 and two later chemistry versions (HiFi-S24, HiFi-N26), at 40x. c. HiFi-N26 against ONT, at 40x. d. Directions of per-read allele length differences between HiFi-R23Q3a and HiFi-R23Q3b at 15,000 random loci. Left, per-locus normalized Mann-Whitney U (Methods): the probability that a randomly chosen HiFi-R23Q3b read carries a longer allele than a randomly chosen HiFi-R23Q3a read. 0.5 (dashed line) indicates no difference. Right, percentage of loci in each direction. P-value, one-sided Wilcoxon signed-rank test against 0.5. e. Alleles detected in HiFi-R23Q3b but absent from HiFi-R23Q3a, cross-tabulated by supporting reads (within 3 bp of the allele length) and length offset from the closest HiFi-R23Q3a allele at the locus. Cell values and shading give allele counts. Offsets begin at 4 bp, the resolution of length clustering (Methods). The same alleles cross-tabulated by difference in motif k-mer count are in Additional file 1: Fig. S5a, and the reciprocal comparison in Fig. S5b.

We next turned to newer HiFi chemistries. We combined the two HiFi-R23Q3 runs, hereafter HiFi-R23Q3, and compared them against two later chemistries at 40x (HiFi-S24, HiFi-N26; n = 1,925,515). The number of loci carrying a replicate-unique allele fell from 57,051 for HiFi-R23Q3 to 41,345 for HiFi-S24 and 30,718 for HiFi-N26 (Fig. 3b). Across all three, 191,941 loci (10%) carried an allele that another dataset lacked, 12,731 (0.7%) carried one supported by at least two reads of the same length, and 62,135 (3%) carried one more than 6 bp from every allele in the other datasets (Additional file 1: Fig. S4c, d).

We then asked how ONT compares. At 1,926,920 loci covered by both, we found ONT carried an allele absent from HiFi-N26 at 457,443 loci (24%), whereas HiFi-N26 carried an allele absent from ONT at 44,106 (2%), a tenfold difference (Fig. 3c). Requiring two-read support gave 43,571 for ONT and 2,596 for HiFi-N26, and requiring an offset of more than 6 bp gave 147,398 and 13,952 (Additional file 1: Fig. S4e, f). ONT thus had significantly more replicate-unique alleles than HiFi.

For our purpose of mutation calling, it was critical whether the discrepancies between HiFi replicates exhibited a consistent directional bias. We thus estimated the probability that a HiFi-R23Q3b read carries a longer allele than a HiFi-R23Q3a read at 15,000 randomly sampled loci (normalized Mann-Whitney U; Methods). Under symmetric error, this value should center on 50%. Instead, we found HiFi-R23Q3b reads longer at 57.5% of loci, and HiFi-R23Q3a reads at 30.5%, with no difference at 12% (Fig. 3d; one-sided Wilcoxon signed-rank test against 0.5, P < 10⁻³⁰⁰). While its cause remains unclear, this non-random discordance limits how two datasets can be compared.

### Thresholds on three read-level features remove nearly all errors between replicates

To find thresholds that can exclude the error set, we took HiFi-R23Qa as the base sample and characterized each HiFi-R23Q3b allele absent from HiFi-R23Q3a by three read-level features: the number of supporting reads, the length offset from the nearest HiFi-R23Q3a allele at the locus, and the change in motif k-mer count relative to that allele (Fig. 3e, Additional file 1: Fig. S5a, and the reciprocal comparison in Additional file 1: Fig. S5b; Methods).

We observed replicate-unique alleles crowded into a narrow part of the space. Of 63,050 alleles differing by ≥3 bp in length (Fig. 3e), 67% were supported by a single read, with counts falling steeply with each additional read. We had already collapsed lengths within 3 bp of an allele the other run carried (Methods), but 58% of the remainder lay within 6 bp and fewer at each larger offset. Change in motif k-mer count discriminated less on its own but flagged cases that point to indels in the flank rather than in the repeat (Additional file 1: Fig. S5a). Interestingly, two of the 63,050 alleles sat well outside this pattern, each with 12 and 18 supporting reads at an offset of 58 and 236 bp (Fig. 3e, top right). We cannot verify whether they are true mutations or errors, so we leave them as an open case.

Two runs of one library a week apart are the most favorable comparison available. We therefore considered the distributions above as a lower bound on error. Real-world analyses are harder: child against parents or a tumor against normal, from separate libraries and sometimes different chemistries. As a result, we repeated the analysis across HiFi-R23Q3, HiFi-S24, and HiFi-N26. We ran TRumut on every ordered pair and chose the smallest thresholds at which the six comparisons averaged fewer than one call each. That criterion gave at least seven supporting reads, a length offset of at least eight bp, and a change of more than three motif k-mers. Rerunning the six ordered comparisons at the chosen thresholds yielded 4 de novo and 3 postzygotic events (Additional file 1: Figs. S6, S7, S8 and S9). We used these thresholds in every analysis below.

The cross-replicate comparison above raised the question of how well existing callers filter out these replicate differences. We ran TRGT-denovo [30] on HiFi-S24 and HiFi-N26, treating one as the parent and the other as the child. Calls were then filtered using the criteria its authors recommend (de novo coverage ≥ 5, allele de novo ratio ≥ 0.7, child de novo ratio 0.3-0.7, no allelic dropout). On such negative control, TRGT-denovo reported 129 de novo loci with HiFi-N26 as the child and 48 with HiFi-S24 as the child. 30 and 21 of the “de novo” calls, respectively, were length changes of 8 bp or longer. In comparison, TRumut only reported one de novo call in each pair.

### De novo tandem repeat mutations across a four-generation pedigree are separable from sequencing error

We applied the TRumut trio module to the CEPH 1463 pedigree, calling each sequenced member against its parents (Methods). We call an allele de novo when neither parent carries it and it occupies the child’s entire haplotype, as a mutation arising in a parental germline or before the first cleavage must (Methods). An allele absent from both parents but confined to a subset of reads on its haplotype is referred to as postzygotic (next section). At chr4:116,297,119-116,297,247, for example, the child carries a parental allele and a second allele more than 20 bp from either parental length on every read of its haplotype (Fig. 4a).

**Figure 4.**
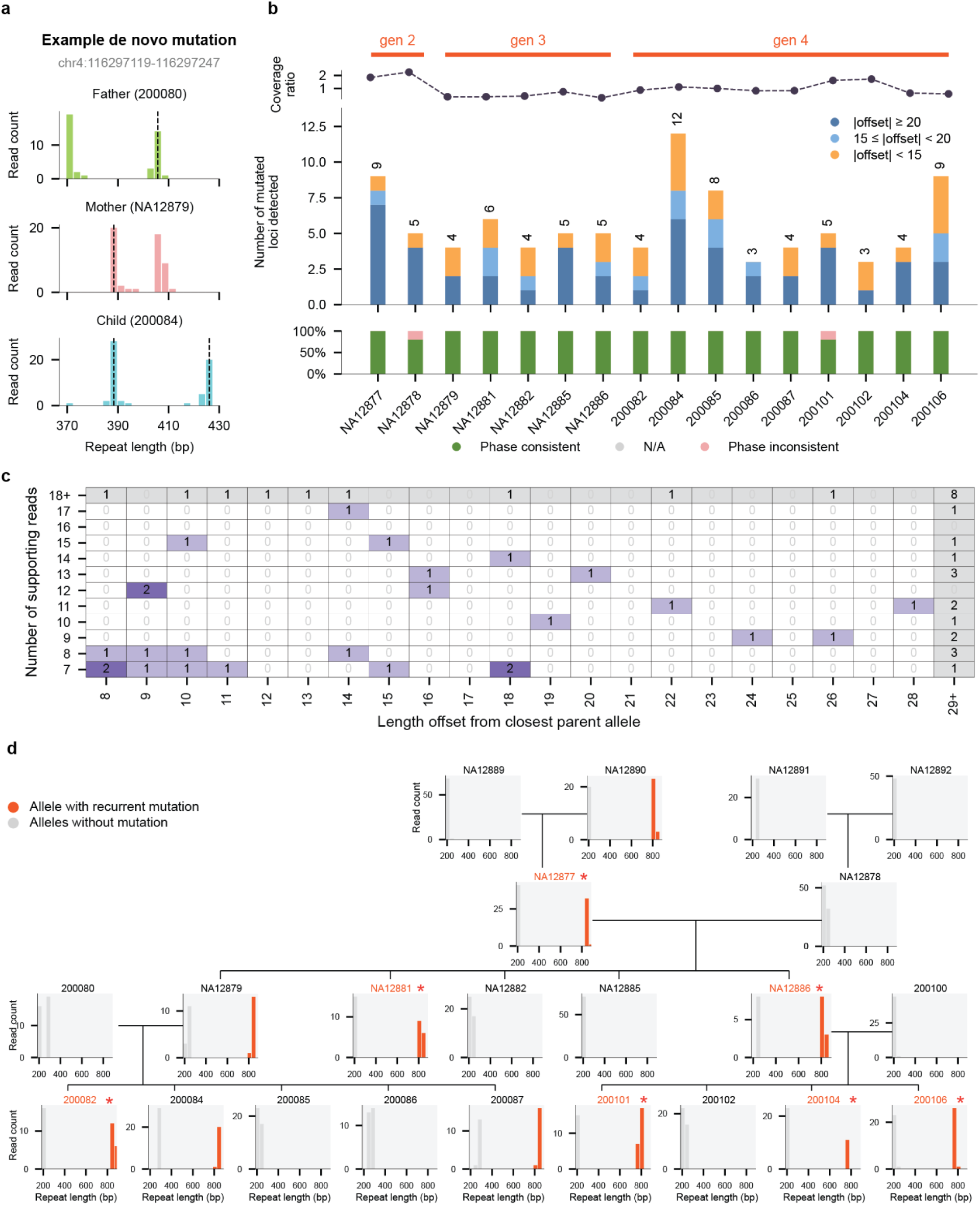
De novo tandem repeat mutations across the CEPH 1463 pedigree. a. Read-level allele lengths at a de novo locus (chr4:116,297,119-116,297,247) in a father, mother, and child. Histograms give reads per allele length. Dashed lines mark the parental alleles transmitted to the child. The child carries an allele absent from both parents on every read of its haplotype. b. Number of loci with de novo mutations in each pedigree member, called against the member’s parents, by generation. Top, coverage relative to the parents’ mean. Middle, number of de novo loci, colored by the length offset of the largest-offset event at each locus from the nearest parental allele. Bottom, proportion of events whose supporting reads derive from one haplotype (phase consistent), from both (phase inconsistent), or from no haplotagged read (N/A). c. De novo events in fourth-generation members (n = 58; a locus with two mutant alleles contributes two events), cross-tabulated by supporting reads and the length offset from the closest parental allele. The final row and column aggregate larger values. The same alleles cross-tabulated by difference in motif k-mer count are in Additional file 1: Fig. S5c. d. Read-level allele lengths at chr4:76,608,528-76,608,747 for each sequenced member of the pedigree, arranged by pedigree position. Red marks the allele lineage in which mutations were called in members of different generations. All other alleles are gray. Asterisks and red labels mark the individuals in which we made a call. Locus filters as in Fig. 3.

Across the 16 members with sequenced parents, we detected de novo mutations at 90 loci, including 14, 24 and 52 in the second, third and fourth generations, respectively (Fig. 4b; by motif length, Additional file 1: Fig. S10). Counts per member were not too different, 3 to 12, and depended only weakly on depth: 9 and 5 in the second generation at twice parental depth, 4 to 6 in the third at half, and 3 to 12 in the fourth at parental depth or above.

We first assessed whether any read from the child’s other haplotype carried a length close to the de novo allele, which a mutation arising on one parental chromosome should not produce. At 88 of 90 loci, the other haplotype carried no such read (Fig. 4b, bottom). The two exceptions were in NA12878 and 200101, the members sequenced above parental depth, where we expected false positives to concentrate.

We also checked how far the calls sat from the thresholds. Among the 58 events at the 52 fourth-generation loci, 37 were offset from the nearest parental allele by 16 bp or more (Methods), and 36 were supported by more than 10 reads (Fig. 4c). When we applied the same caller to HiFi datasets of HG002, where no true de novo alleles are expected, four of six comparisons had yielded no calls and the other two yielded two each (Additional file 1: Figs. S6 and S7). We found the few calls that passed through the thresholds sat close at the decision boundaries, with all four supported by eight or fewer reads.

A real de novo allele should be inherited, so we used the next generation to test the calls. Four members with de novo calls had sequenced children, giving 23 alleles and 110 transmissions (Additional file 2: Table S1). For each transmission, we classified the allele the child received from the carrier parent as the de novo allele, the parent’s other allele, or neither (Methods). 20 of the 23 alleles were transmitted to at least one child. Across 110 individual transmissions, the de novo allele was passed on in 43 and the parent’s other allele in 50. This near-equal transmission ratio is consistent with mendelian segregation of true de novo mutations. In the remaining 17 pairs, the child carried a length matching neither. We suspect that such cases reflect secondary mutations at highly mutable loci. Interestingly, among these calls, one allele at chr4:76,608,528-76,608,747 changed length in seven members across generations, expanding in some transmissions and contracting in others (Fig. 4d; Additional file 1: Fig. S13 shows a second example). It is a reoccurring de novo mutation.

We next benchmarked our results against the de novo tandem repeat calls that Porubsky et al. [31] generated, for the same pedigree, using TRGT-denovo [30] (Additional file 1: Fig. S11). To make the two sets comparable, we re-ran TRumut on their chosen catalog and compared the five third-generation members present in both sets (Methods). Homopolymer loci were excluded as elsewhere. We found that 24 of our 25 de novo calls were in the published set as well. The one call not included in the set is an 821 bp allele in NA12881 at a GCATA repeat. Nine reads on a single haplotype support it, all at least 10 bp shorter than every paternal read (831 to 874 bp) and with a matching loss of motif k-mers. We therefore consider it to be a paternal contraction the published pipeline missed.

The larger question was why 348 out of 372 published loci were not called with our method. To begin with, 248 loci had published size changes of 7 bp or less, which our replicate analysis showed to be indistinguishable from sequencing error. Porubsky et al. list 79% of such loci in NA12879 and NA12886 as transmitted to at least one G4 child, somewhat below the 87% (20 of 23) we observe from our own calls. When we evaluated the other 100 against our read-level filters, 61 failed at least one of them: 26 were within 8 bp of a parental read, 33 had fewer than seven supporting reads, and two had insufficient change in motif k-mer count. Another 16 were at loci where the child or a parent had fewer than 10 reads, and five we called but did not classify as de novo, for instance, because the reads containing them were unphased. In short, every difference so far came down to a threshold specific to our method.

The remaining 18 loci all fell in regions of poor mappability (Additional file 3: Table S2): six were subtelomeric, two pericentromeric, eight within other annotated segmental duplications, and two in the pseudoautosomal region. The flanking uniqmers at these loci were 44 kb to 20 Mb apart or missing on one side, and TRumut therefore made no call. Alignment, however, still produced calls at all 18. Six of these were flagged by Porubsky et al. themselves as recurrent. When we looked at the other 12 that passed their filters, eight were supported by reads with a median mapping quality below 10. Even at loci with median MAPQ 60, the ambiguity of the repeat sequence led to soft clipping and inconsistent alignments (Fig. S12). Since reads in these regions cannot be placed or aligned unambiguously, we consider the absence of a call to be the right answer here. In summary, we conclude that the difference between the two call sets largely reflects our error-based filtering rather than missed mutations.

We also compared the runtime and peak memory of the two pipelines. On the same 22 genomes with 8 CPUs per run, TRumut allele extraction used eightfold less CPU time than TRGT genotyping (median 1.6 vs 13.1 CPU-hours per genome) and sixfold less peak memory (3.1 vs 17 GB). Calling de novo mutations in a trio end to end cost us 12-fold less CPU time (4.5 vs 55.5 CPU-hours, median of 16 trios; Methods). TRumut therefore runs an order of magnitude cheaper.

### Somatic tandem repeat mutations can be called above the error floor, at the cost of sensitivity

Postzygotic (or somatic) mutations arise after the first cleavage in the embryo and are therefore carried by a subset of cells (Fig. 5a). We called postzygotic mutations at 56 loci across the 16 members with sequenced parents. The second, third, and fourth generations contributed 21, 11, and 24, and two members carried none (Fig. 5b). Counts were lower than for de novo events and more uneven: NA12877, sequenced to roughly twice parental depth, alone carried 17, and no other member carried more than six.

**Figure 5.**
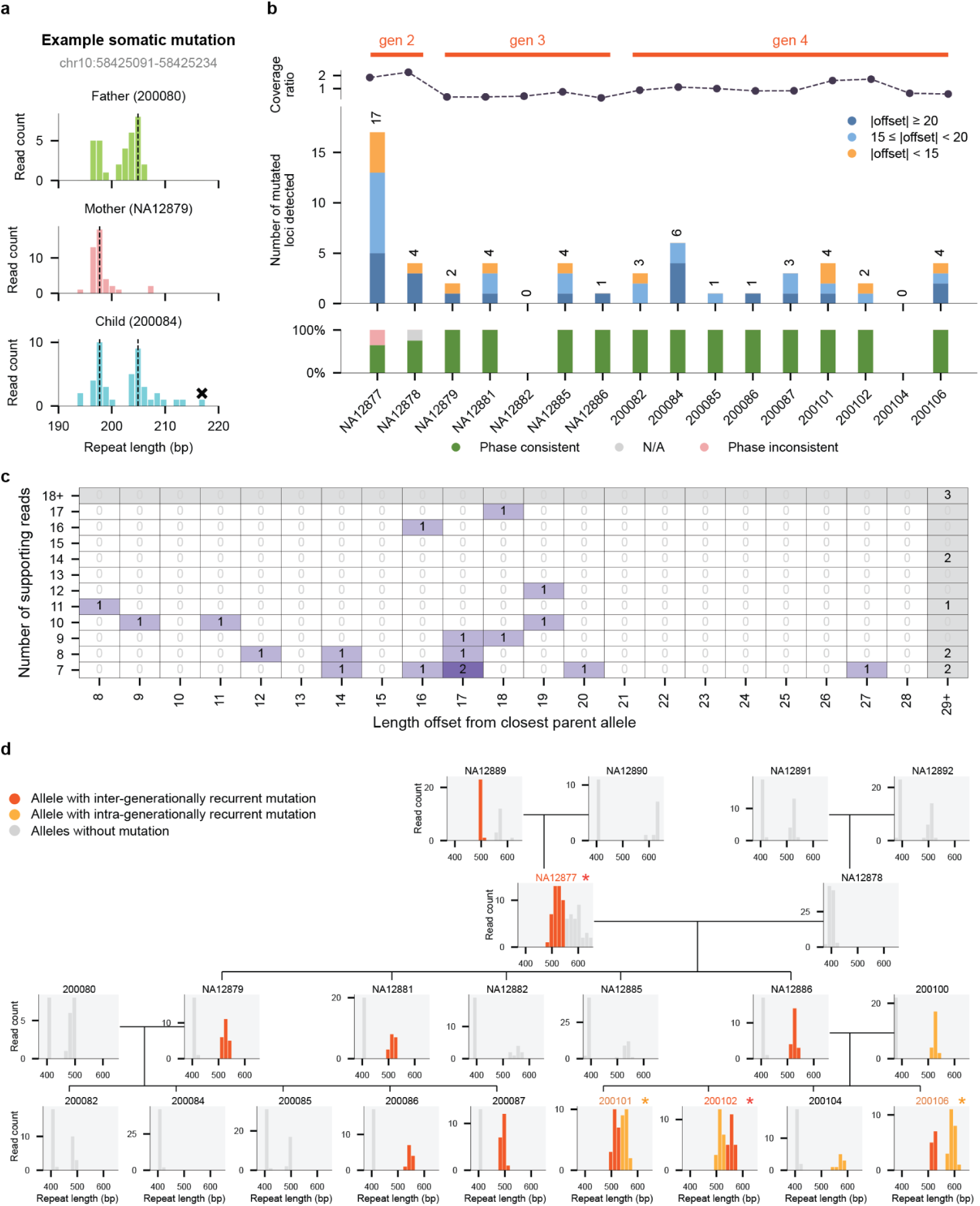
Postzygotic tandem repeat mutations across the CEPH 1463 pedigree. a. Read-level allele lengths at a postzygotic locus (chr10:58,425,091-58,425,234) in a father, mother, and child. Histograms give reads per allele length. Dashed lines mark the parental alleles transmitted to the child. The child carries an additional allele on a subset of reads (marked x), the signature of a postzygotic event. b. Number of loci with postzygotic mutations in each pedigree member, called against the member’s parents, by generation. Top, coverage relative to the parents’ mean. Middle, number of postzygotic loci, colored by length offset of the largest-offset event at each locus from the nearest parental allele. Bottom, proportion of events by phase consistency. c. Postzygotic events in fourth-generation members, cross-tabulated by supporting reads and the length offset from the closest parental allele. The final row and column aggregate larger values. The same alleles cross-tabulated by difference in motif k-mer count are in Additional file 1: Fig. S5d. d. Read-level allele lengths at chr11:90,271,112-90,271,289 for each sequenced member, arranged by pedigree position. Color marks one lineage tracked through the pedigree: red, mutations called in members of different generations; orange, mutations called in members of the same generation; gray, alleles with no call. Asterisks and colored labels mark members with a call. Locus filters as in Fig. 3.

We expected postzygotic calls to sit closer to the read-support threshold than de novo calls, since a somatic allele is present in only a fraction of reads by definition. Indeed, among the 24 fourth-generation calls, 15 were supported by fewer than 10 reads (Fig. 5c). Offsets, by contrast, were not small. Most lay 15 bp or more from the nearest parental allele, and about half 20 bp or more. Overall, 2.7 postzygotic calls were made per fourth-generation member, against the 0.5 average in HG002 replicates (Additional file 1: Figs. S8, S9). We therefore acknowledge that some postzygotic calls may reflect persistent errors, but most demonstrate signal strength exceeding errors.

Individual loci further supported our results. At chr11:90,271,112-90,271,289, the lineage that acquired transmitted changes across generations also acquired somatic changes within individuals, while other alleles at the locus acquired neither (Fig. 5d). Error would not select one specific allele and then recur in member after member, so we consider these postzygotic mutations to be genuine.

### Cancer cell lines differ in repeat mutation clonality but share recurrent loci

We applied the TRumut cancer module to six HiFi tumor-normal pairs (COLO829, HCC1395, HCC1937, HCC1954, H1437, and H2009; Methods), reporting tumor alleles absent from the normal under the thresholds above. We classified each event by the reads carrying it (Fig. 6a; Methods). We call an event clonal when the mutant length is the only peak on its haplotype, subclonal when it forms a smaller peak alongside the unmutated allele, and private when it is carried by too few reads to form a peak at all.

**Figure 6.**
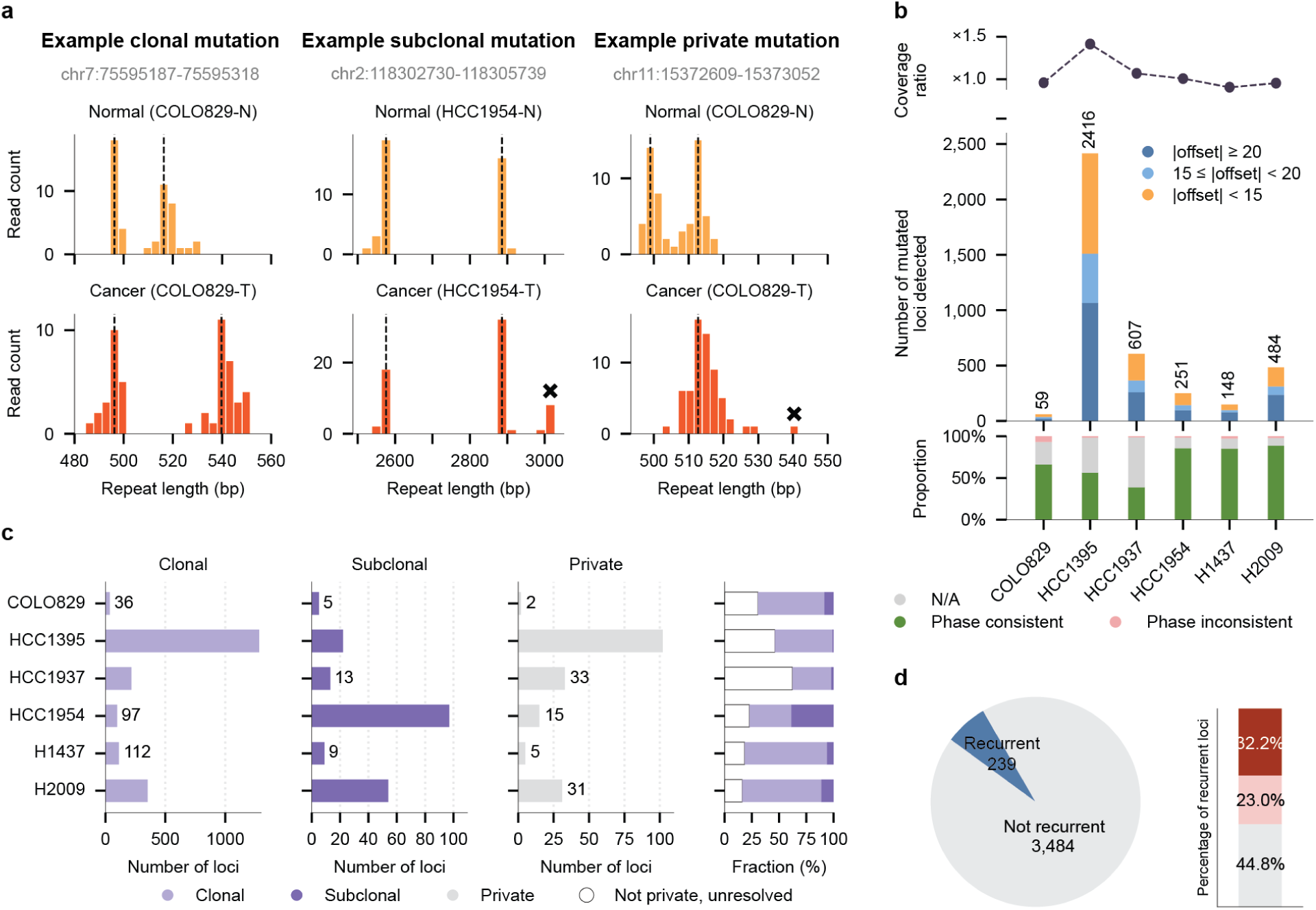
Somatic tandem repeat mutations in six cancer cell lines. a. Read-level allele lengths at three loci, one per event class, in a tumor and its matched normal. Histograms give reads per allele length, and dashed lines mark alleles in the normal. Left to right, a clonal, subclonal (x), and private (x) mutation, as defined in the text. b. Number of mutated loci per cell line. Top, tumor coverage relative to the matched normal. Middle, number of loci, colored by length offset of the largest-offset event at each locus from the nearest normal allele. Bottom, proportion of events by phase consistency. c. Class composition of mutated loci. Left, number of loci called clonal, subclonal, and private per line. Right, the same as proportions, including loci unclassifiable without phase (not private, unresolved; Methods). d. Left, recurrence across the six cell lines. Right, concordance of events at recurrent loci. Events match in direction when all are expansions or all contractions, and in length when the mutant allele lengths also agree within 7 bp (Methods). Locus filters as in Fig. 3.

We found mutated loci ranged from 59 in COLO829 to 2,416 in HCC1395 (Fig. 6b). 38% of these calls lack phasing information primarily due to loss of heterozygosity in cancer genomes. Of the remaining calls that can be phased, 97% are phase-consistent, implying high accuracy (Fig. 6b, bottom).

We next categorized somatic events as clonal, subclonal, or private to resolve their evolutionary timing in the tumor (Fig. 6c). Clonal events formed the largest resolved class in every line except HCC1954. There, subclonal events matched clonal ones at 97 each, roughly two to 20 times the subclonal count in any other line (Additional file 1: Fig. S14). HCC1954 has about 90 chromosomes. The high fraction of subclonal calls implies that many somatic mutations in tandem repeats arose after chromosomal duplications. Subclonal events were otherwise few, from five in COLO829 to 54 in H2009. As expected of our stringent read support thresholds, only a minority of calls were private, from two in COLO829 to 102 in HCC1395. Because unphased reads left a substantial share of non-private events unclassified, so we regard clonal and subclonal counts as lower bounds.

Lastly, we asked whether the same loci mutated in more than one line (Fig. 6d). Of the 3,723 loci mutated in at least one line, 239 were mutated in two or more. This result strictly outperformed 10,000 permutations from a conservative null model, where we randomly assigned per-line events across mutated loci (null mean 140; P < 10⁻⁴; Additional file 1: Fig. S15a). At the recurrent loci, mutations in different lines matched in length in 32% of cases, matched in direction but not length in 23%, and differed in direction in 45%. Some loci are therefore mutable in more than one tumor.

### Large length changes recur at shared loci and are more often clonal in tumors

Length changes of 100 bp or more are the class most implicated in disease [16]. We called them with the offset threshold raised to 100 bp and no minimum read support, since replicate error is sparse at such offsets (Methods; Additional file 1: Fig. S15). Between HG002 datasets, this gave 62 on average (Fig. 7a).

**Figure 7.**
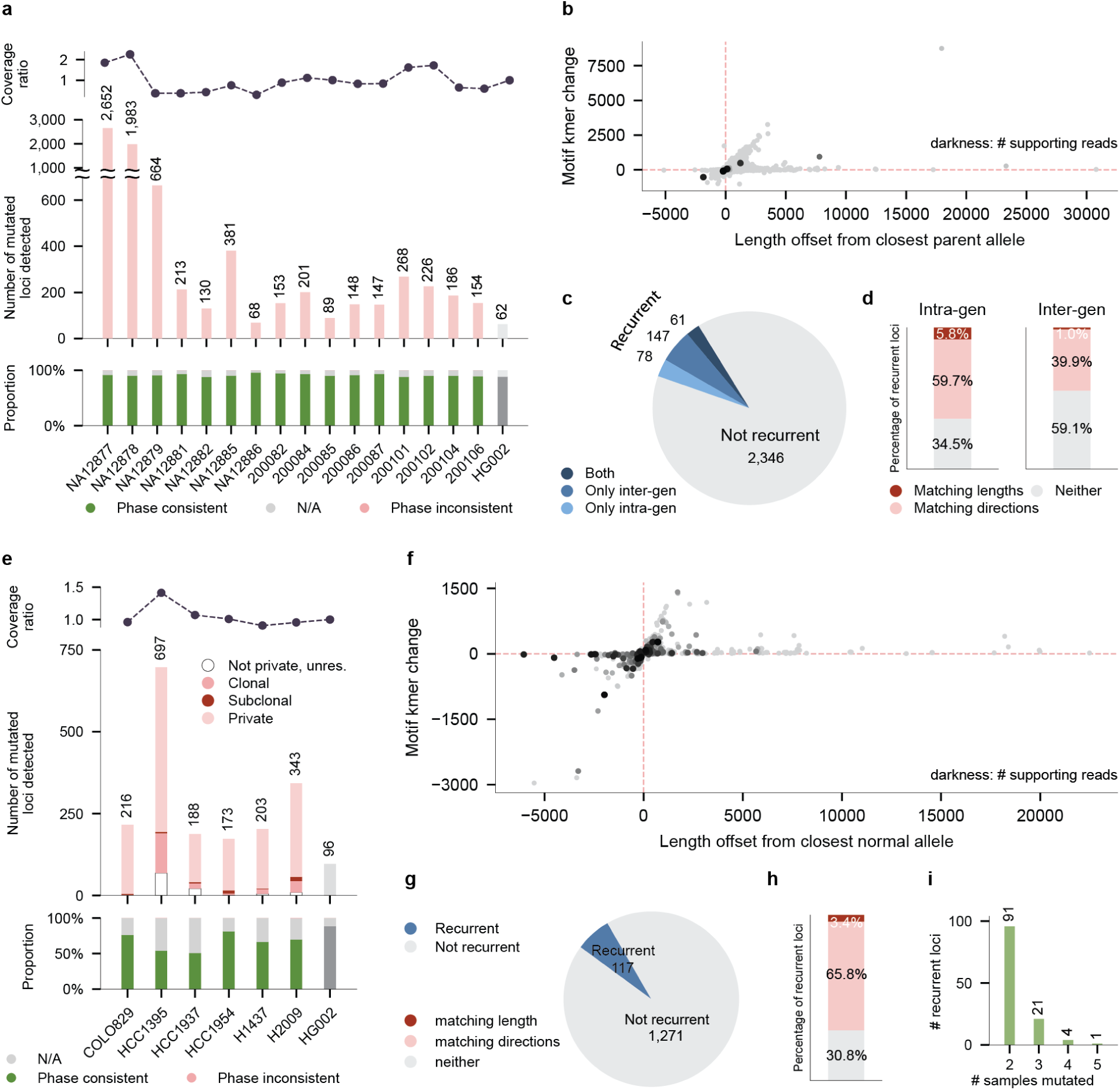
Large length changes in tandem repeats across the pedigree and cancer cell lines. Events are large length changes, at least 100 bp in either direction from the nearest allele in the comparison sample (parents or matched normal). Locus filters as in Fig. 3. a. Events per pedigree member, called against the member’s parents. Top, coverage relative to the parents’ mean. For HG002, coverage is the test dataset relative to the comparison dataset. Middle, number of loci with a call. Bottom, proportion of loci by phase consistency. The rightmost bar (HG002) is a negative control, the mean over four comparisons between HG002 HiFi datasets (HiFi-R23Q3a and HiFi-R23Q3b in both directions, HiFi-S24 and HiFi-N26 in both directions). b. Length offset from the closest parental allele against the change in motif k-mer count, one point per event across all members. Darkness gives supporting reads, and dashed lines mark zero. HG002 comparisons excluded. c. Recurrence among third- and fourth-generation members, whose depths are comparable. We classify recurrent loci as intra-generational, inter-generational, or both (Methods). d. Concordance at recurrent loci, defined as in Fig. 6d, separately for intra- and inter-generational recurrence. e. Events per cancer cell line, called against the matched normal. Top, tumor coverage relative to the matched normal. Middle, number of loci colored by event class, including loci unclassifiable without phase (not private, unresolved; Methods). Bottom, proportion of loci by phase consistency. The rightmost bar (HG002) is the same negative control as in a, the mean over four comparisons between HG002 HiFi datasets, restricted to the same loci as the cell lines. f. As in b, for the cancer cell lines, with offsets measured from the closest normal allele. HG002 comparisons excluded. g. Recurrence across the six cell lines. h. Concordance of events at recurrent loci, defined as in d. i. Number of recurrent loci by the number of cell lines carrying an event.

In the pedigree, the number of loci that underwent large length changes ranged from 68 to 2,652. Most of the events we found came with a proportional change in motif k-mer count (Fig. 7b). Nearly all events, 99%, are supported by a single read. We therefore expected counts to be driven by depth, and they indeed tracked coverage relative to parents. The two over-sequenced second-generation members had 2,652 and 1,983 while others ranged from 68 to 664. Notably, this excess did not disappear when we downsampled all members to similar coverage nor did it track age at sampling. The source of this coverage-independent excess therefore remains unresolved.

We then checked whether the same loci mutated in more than one member. Among the third- and fourth-generation members, whose depths are comparable, 286 of 2,632 mutated loci were recurrent: 61 within a generation, 147 across generations, and 78 both (Fig. 7c). This exceeds every one of 10,000 permutations under the same null as before (mean 264; P < 10⁻⁴; Additional file 1: Fig. S15b). Large length changes in the pedigree therefore recur at specific loci, although the excess over chance is modest. At the recurrent loci, events matched in direction at 66% of intra-generational and 41% of inter-generational loci, and in length at only 6% and 1% (Fig. 7d). Note that the permutation test only suggests large changes are correlated and does not explain what caused the correlation.

The cancer lines differed from the pedigree in both the clonality and the direction of large length changes. Of the 173 to 697 changes per cell line, we classified 19% as clonal or subclonal (Fig. 7e). In the CEPH pedigree, by contrast, only 0.4% were (Fig. 7a). We also observed a shift towards contractions, which accounted for 67% of tumor events but 49% of pedigree events (Fig. 7b, f). We thus conclude that large changes in tumors more often reach a substantial fraction of cells, and more often shorten the locus, than those in healthy individuals.

Recurrence across cancer cell lines, on the other hand, looked much the same. Of 1,388 loci mutated in any line, 117 were recurrent, 26 of them in three or more lines (Fig. 7g-i). As in the pedigree, the count exceeded every permutation under the null (mean 95; P < 10⁻⁴; Additional file 1: Fig. S15c). At the recurrent loci, we saw events matched in direction 69% of the time, higher than in either pedigree comparison, and in length only 3% of the time (Fig. 7h). Large changes thus recur at specific loci in both settings, but reach a substantial fraction of cells only in tumors.

### A population panel extends calling to samples with no matched control

The trio and cancer modules identify mutations by absence from a comparison sample. Archival tumors, many cell lines, and most individuals have no such comparator. In its place, we built a panel of normals from the reads of 232 HPRC genomes (Fig. 8a) [32]. At each locus, the panel records the range of allele lengths in the population, excluding rare lengths (Methods). With the panel module (`trumut use-pon`), we report test-sample alleles outside that range under the same thresholds as the other modules.

**Figure 8.**
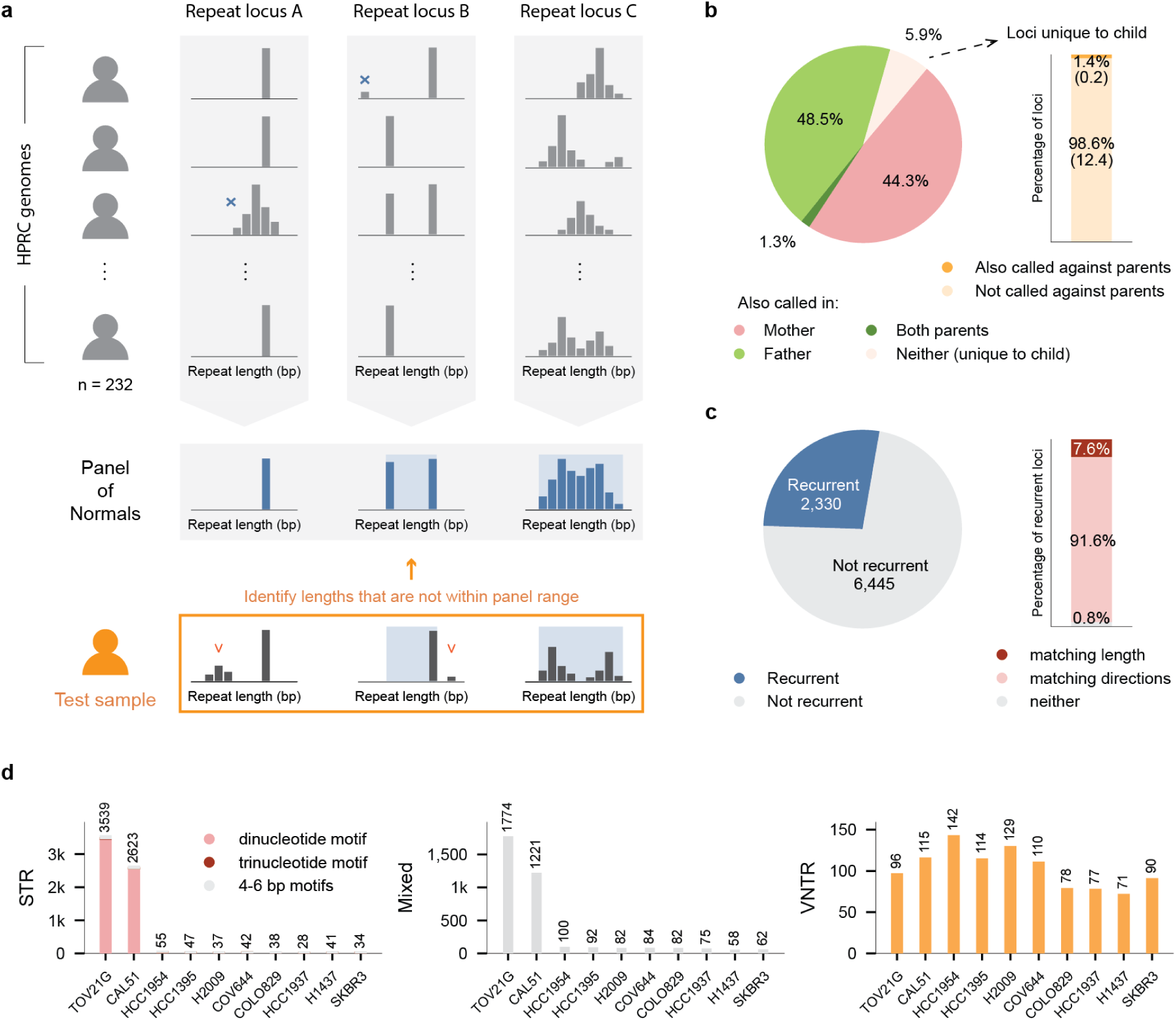
Calling extreme tandem repeat alleles against a panel of normals. a. Schematic of panel-based calling. At each locus, allele lengths across 232 HPRC genomes (top) define the population range, drawn as a blue band (middle). Rare lengths (Methods) are excluded and marked with a blue cross (top). In a test sample (bottom), reads outside the band are evidence of an extreme allele, and reads inside it are set aside. b. Origin of panel-based calls in each child, averaged across children. Left, proportion of a child’s calls also called against the panel in the mother, father, both, or neither. Right, proportion of child-unique calls also recovered by the trio module. Values in parentheses, mean loci per child. c. Left, recurrence across cancer cell lines. Right, concordance at recurrent loci, as in Fig. 6d (Methods). d. Number of extreme alleles per cancer cell line, called against the panel, by repeat class: STR (left), mixed (middle), VNTR (right). STR bars are stacked by motif length: dinucleotide, trinucleotide, or 4-6 bp. Locus filters as in Fig. 3.

A panel call, however, flags an allele as extreme, lying outside the lengths seen in the population, while a trio call flags it as new in the individual. An allele can be either, both, or neither, and we expect the alleles that matter most biologically to be both. To see how the two calls relate, we reasoned as follows. A panel call in the child is (i) extreme but not new if a parent shares it and it is clonal in the child, (ii) extreme and new if no parent has an equivalent call, or (iii) extreme and new regardless of parental calls if it is subclonal or private in the child.

We called every CEPH 1463 member with sequenced parents (n = 16) against both the panel and its parents (Additional file 1: Fig. S17). Almost every panel call in a child was also a panel call in a parent: 49% in the mother, 44% in the father, and 1% in both (Fig. 8b). Of the shared calls, 94% were clonal in the child and thus not new (i), and 2% were subclonal or private (iii), the child having mutated postzygotically where a parent already carried an extreme allele. Only 6% of panel calls were unique to the child (ii), and few of them also passed the trio module’s thresholds (1% of them, 0.2 loci per child on average). When we looked from the trio side, the overlap was just as small: most trio calls were not panel calls, indicating most mutant alleles fell within the population range. We conclude that each module finds mostly alleles the other does not. Inherited extreme alleles need the panel, new alleles within the population range need a comparator, and alleles both extreme and new, the smallest set here and the one most worth finding, require the two together.

We next extended the panel module to 10 cancer cell lines. Of 8,775 loci with an extreme allele called in any line, we found 2,330 in two or more lines, far exceeding the permutation null (Additional file 1: Fig. S18). Furthermore, 99.2% of extreme alleles at recurrent loci matched in direction, with 8% matching even the length (Fig. 8c).

The panel also let us compare microsatellite-stable and unstable lines (Fig. 8d). We found the two microsatellite-unstable lines, TOV21G and CAL51, carried roughly 60- to 90-fold more extreme alleles in STRs than the other eight lines (3,539 and 2,623 versus 28 to 55) and twenty times more in mixed-motif loci (1,774 and 1,221 versus 58 to 100). In VNTRs, their extreme allele counts were 96 and 115, falling within the 71 to 142 of the other lines. Microsatellite instability therefore destabilizes short-unit and mixed-motif repeats but not VNTRs.

### Large length changes can be called from a single sample as outlier alleles

The panel compares a sample with the population, so an allele needs only to clear the usual offset threshold (8 bp) beyond the population range to be called, and it may be inherited, de novo, or somatic. The alleles that matter most in repeat expansion disorders are different in two ways. They are large, often hundreds of base pairs [16], and they are often mosaic, present in a minority of cells and therefore on a few reads [5,8]. We asked whether such alleles could be found from a sample alone, with no comparator at all. In the HG002 replicates, an allele 100 bp or more from every other allele at its locus rarely arose from error, even on a single read (Fig. 3e; Additional file 1: Fig. S16). We therefore term an “outlier” the longest or shortest allele at its locus, at least 100 bp from any other allele in the same sample, with no read-support minimum (Methods). Because a germline allele occupies its whole haplotype, an outlier on a small fraction of reads can only be somatic or error, and the replicate baseline bounds the error.

Across the 10 cell lines, we found coverage-normalized outlier counts from 3.7 to 14.4, against a mean of 4.48 across the HG002 datasets, our negative control (Fig. 9a). Three lines exceeded HG002 by more than two-fold: H2009, COV644, and SKBR3. Interestingly, we saw no dependence on microsatellite instability in any repeat class (Fig. 9b). The two MSI lines carried 11 to 18 STR outliers against 3 to 47 in the rest, as opposed to the 60- to 90-fold STR excess seen under the panel (Fig. 8d). We interpret this result as mismatch repair loss producing many small slippage events but not large ones. Lines ranked consistently across classes, COV644 highest in all three. We also checked that the criterion does not simply select the longest loci: outlier loci had a median catalog length of 562 bp, with 74% under 1 kb. Offsets split 43% expansions to 57% contractions (Fig. 9c).

**Figure 9.**
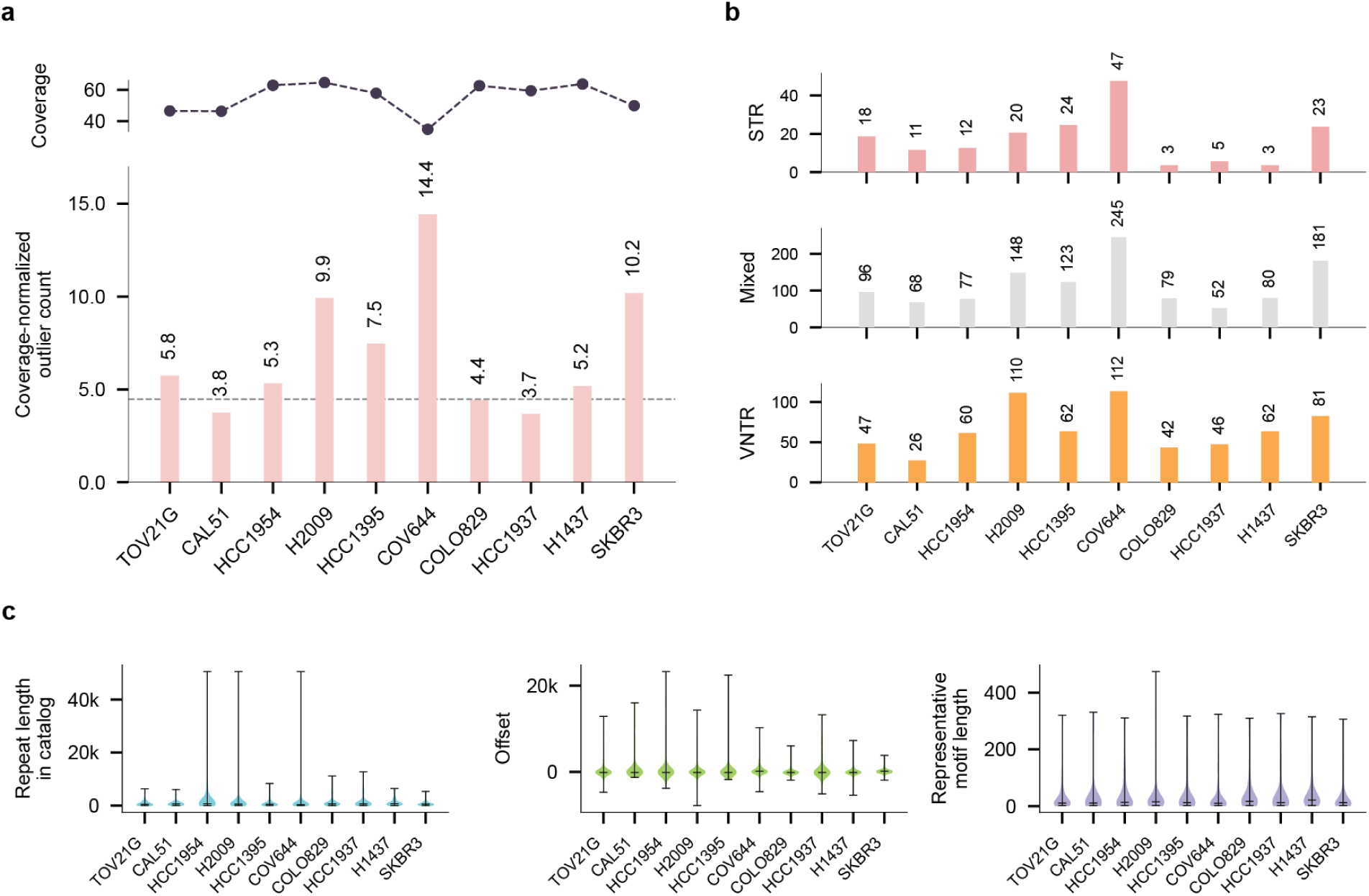
Outlier tandem repeat alleles across cancer cell lines. An outlier allele is the longest or shortest allele at its locus, at least 100 bp from the next. Homopolymers were excluded (Methods). a. Outlier alleles per cell line. Top, coverage. Bottom, outlier count normalized by coverage. Dashed line, negative control: the mean coverage-normalized outlier count across three HG002 HiFi datasets (HiFi-R23Q3, HiFi-S24, and HiFi-N26). b. The same alleles by repeat class: STR (top), mixed (middle), VNTR (bottom), not coverage-normalized. c. Outlier properties per cell line. Left, reference length of the repeat. Middle, offset to the next-closest allele. Right, length of the motif spanning the most reference sequence (Methods). All values are in base pairs.

## Discussion

Most of what we know about somatic tandem repeat mutations comes from summary statistics rather than individual mutations. Callers report per-locus instability scores [27], stutter parameters [33], or an aggregate burden per sample [25,26]. We argue that this is due to representational constraints rather than a lack of biological interest. Specifically, a tandem repeat allele is represented in standard pipelines as a count of repeat units within a predefined reference interval. Alignment ambiguity near repetitive regions, however, causes these interval boundaries to fluctuate across reads. This forces pipelines to report only the summary statistics. We argue that this has held back our understanding of how repeats mutate in somatic cells, which matters for development, aging, tumor evolution, and repeat expansion disorders.

We demonstrate here that comparing individual alleles, and therefore calling mutations, becomes possible once the allele is reliably represented the same way on every read. Our method takes the sequence between two pangenome-conserved anchors and describes it by raw sequence, length, and motif k-mer content. By bypassing sequence alignment, we were able to define alleles consistently across reads.

The practical consequence is that error becomes measurable. We found that roughly one locus in ten disagrees at the read level, and one in 150 even with two concordant reads. To our knowledge, this represents the first controlled error benchmark for tandem repeat in long-read sequencing. This result concerns the data type, not a specific caller, and thus constrains every long-read study of somatic tandem repeat mutation. It is also invisible to current benchmarks. Standard comparisons are made between consensus genotypes, which averages away exactly the variation that exposes sequencing error.

Informed by this error analysis, we derived filtering thresholds to be used in standard mutation calling practices. The resulting tool, TRumut, represents the first method for tandem repeats with capability to resolve both postzygotic and somatic mutations from long reads.

In particular, our results in cancer point to two things worth pursuing. First, we saw somatic mutations recur at shared loci, including kilobase-scale VNTRs that short-read surveys of recurrent expansion could not reach [10]. Whether these loci are selected or simply mutable in a shared cellular context is an open question. Second, we found that microsatellite instability increased extreme STR alleles by two orders of magnitude while leaving VNTRs unchanged. The same pattern was seen in yeast where loss of mismatch repair destabilized short-unit repeats and spared long ones [34]. It could not be tested in human tumors though, because short-read surveys of MSI could not measure VNTRs [35,36]. Our data reaffirm the yeast result. They also attest to how little we know about long-repeat instability in cancer.

Some limitations follow from how we call mutations. We compare allele length and motif k-mer content between samples, so a mutation that rearranges internal sequence while leaving both unchanged is not reported. We also cannot call the 0.3% of catalog repeats that have no anchor pair, though most are near telomeres and centromeres. Other limitations concern interpretation. Our conservative thresholds discard true events that are indistinguishable from error. As in any somatic analysis, we also found that sensitivity scaled with coverage. For these reasons, we do not regard the counts we obtain as mutation rates.

## Conclusions

Alignment and sequencing error, rather than read length, have limited long-read studies of somatic tandem repeat mutations. By calling read-level alleles without alignment, we measured this error for the first time and derived thresholds to remove it. With those calls we saw four things. In a four-generation pedigree, one inherited allele mutated repeatedly along a lineage while its homolog did not. In cancer cell lines, somatic mutations recurred at shared loci. Large length changes recurred in the pedigree as well, but reached a substantial fraction of cells only in tumors. Against a population panel, we saw microsatellite instability destabilize STRs but not VNTRs, a pattern now measurable in human tumors. Somatic repeat mutations can therefore be separated from error and studied as individual events. We expect this method to let us ask the questions long asked of other somatic mutation types, beginning with when they arise, with what signatures, and under what selection.

## Methods

### Overview of TRumut

TRumut determines the tandem repeat allele carried by each read and uses these read-level data to call tandem repeat mutations in three settings: de novo and postzygotic mutations in trios, somatic mutations in tumors with a matched normal, and extreme alleles against a panel of normals in samples with neither. All analyses used TRumut v0.1.0 with default parameters unless stated. Commands are given in the relevant sections with placeholders in angle brackets. The catalog and panel files are those deposited at Zenodo [37].

TRumut accepts HiFi reads in any common format (SAM/BAM/CRAM/FASTA/FASTQ). A genome-wide repeat catalog is annotated with uniqmers and used in place of standard mapping (see “Selection of anchor k-mers” and “Constructing the tandem repeat catalog”). Allele sequences are read directly from individual reads, and the resulting per-read allele table is the common input to three downstream callers.

### TRumut allele

#### Selection of uniqmers

TRumut locates loci through fixed flanking anchors that represent a single genomic position and rarely mutate. To identify them, we counted k-mers in the 579 haplotype-resolved assemblies of the Open Human Genome Library v4.0 [29] (see Availability of data and materials). K-mers that occurred exactly once in at least 98% of assemblies were retained. We also removed k-mers that match elsewhere in the genome within an edit distance of 3, since a few errors or variants could shift the anchor to that position. We set k = 31, long enough that most of the human genome is single-copy yet short enough that a k-mer rarely spans a variant or error. At this length, 34% of k-mers in the human genome (Additional file 1: Fig. S3a) qualified as uniqmers.

#### Constructing the tandem repeat catalog

The base catalog of tandem repeats were built by running Tandem Repeats Finder [38] on the GRCh38 reference genome. Each entry records the coordinates of one repeat with its motif sequence and length. Repeats longer than 15 kb, beyond typical HiFi read length, were dropped.

We next assigned each repeat its “flanking” uniqmer pair, defined as the last uniqmer ending before the repetitive sequence and the first beginning after it. Uniqmer positions on GRCh38 were obtained with kmer-map (https://github.com/lh3/kmer-map). The median distance between a repeat boundary and the flanking uniqmer was 26 bp (Additional file 1: Fig. S3b). When neighboring repeats overlap, lie within 31 bp of one another, or have no uniqmer in the gap between them, a single anchor pair is assigned. This was unnecessary in 73% of regions, and the maximum number of repeats sharing an anchor pair was 10 (Additional file 1: Fig. S3c). For such regions, we define the representative motif as the constituent motif spanning the most reference sequence. Repeats with no flanking uniqmer, mostly near telomeres or centromeres, were 0.3% of the catalog and were dropped.

Loci on chrY and homopolymers are in the catalog but excluded from every analysis above, including the error characterization. Homopolymers were set aside because it is a known region of high HiFi indel errors, and chrY because it is hemizygous and carries very low uniqmer density (Additional file 1: Fig. S3a).

#### Calling repeat alleles from individual reads

Given these anchors, the allele-calling module scans each read for exact 31-bp matches to the anchor pairs on either strand and extracts the sequence between them. Within this interval, we do not assign repeat boundaries. Instead, we represent each allele as the full inter-anchor sequence and its length. Downstream modules compare these directly across reads and samples.

Because the interval also contains non-repetitive sequence, an indel there may change the inter-anchor length without a repeat mutation present. Downstream modules thus compare motif k-mer counts alongside lengths (see “Motif k-mer counting”).

Anchor matching can still fail on a read through rare variants absent from the HPRC assemblies, errors within the anchor, or structural events. A read is therefore discarded for a locus when either anchor matches zero or multiple positions, or when the two anchors match in an inconsistent relative order, orientation, or strand combination, including overlapping matches. In HiFi-R23Q3a, these filters together left 0.01% of loci with no calls. For aligned input, we also exclude secondary and supplementary alignments.

Per-read allele tables were generated from each haplotagged alignment with trumut allele <sample.bam> <catalog.tsv> <sample.alleles.tsv> --input-type reads, the same command for every dataset in this study.

#### Motif k-mer counting

To assess how much of an inter-anchor sequence is repeat-derived, the module counts motif k-mers in it. A motif k-mer is defined as any k-mer that can arise from tandem repetition of the locus motif. We use k = 7, long enough that matches to unrelated sequences are rare, yet short enough that the imperfections common in real repeats do not break most matches. For example, motif 7-mers for a repeat with “CAG” motif would be CAGCAGC, AGCAGCA, and GCAGCAG. A non-motif k-mer is any other k-mer in the sequence. K-mers are counted in strand-collapsed canonical form. For an anchor pair that merged several repeats, we pool all constituent motifs to create the motif k-mer set.

Motif k-mer counting is a coarse screen rather than an exact measure, since imperfect repeats do not match their motif k-mers exactly. Its role is to screen out false positives driven by non-repetitive sequences.

#### Localizing non-motif indels relative to the repeat

K-mer counting also indicates whether a non-repetitive insertion lies inside the repeat or in its flanks. For an insertion to have been inside a repeat, three conditions must hold: (i) there is an increase in non-motif k-mer counts compared to the comparison sample (the matched normal for cancer, the parents for trios), (ii) a run of consecutive non-motif k-mers exists between two motif k-mers, and (iii) those found consecutively must be among the k-mers whose counts increased compared to the comparison sample. In practice, an interval contributes when it contains at least 10 non-motif k-mers and at least 95% of them occur more often than in the comparator. The reported value is the number of such excess k-mers summed over qualifying intervals.

### Error profiling of HiFi sequencing in tandem repeats

#### Symmetric comparison with TRumut mutual

Replicates are compared against one another rather than against a designated reference, since every replicate carries errors and none is the ground truth. The `trumut mutual` module uses each input in turn as the test sample and each other input in turn as its comparison. At an anchored locus, a test allele length counts as “deviating” when it differs from every length in the comparison set by at least the minimum offset. Deviating lengths are then collapsed, taking the one with the most supporting reads first and absorbing any length within the merge width of it until each has been reported or absorbed. The merge width defaults to 3 bp and the minimum offset to one more, 4 bp. This was selected to be small enough that only near-identical lengths merge and larger differences between replicates survive for the error analysis. For each reported allele, we record its supporting read count, the inputs that lack it, and the signed distance to the nearest allele length in each. Replicates were compared with `trumut mutual <out.tsv> -i <A.alleles.tsv> -i <B.alleles.tsv> [-i <C.alleles.tsv>]`.

#### Sequencing data

We used six sequencing datasets, all of the same Genome in a Bottle sample, HG002. HiFi-R23Q1 is the earliest, from a PacBio Revio run on the platform’s initial chemistry. HiFi-R23Q3a and HiFi-R23Q3b are two replicates from the GIAB PacBio Revio release of 31 October 2023. HiFi-R23Q3 denotes the two combined. HiFi-S24 was generated on a PacBio Revio instrument with SPRQ chemistry, and HiFi-N26 on a PacBio Revio instrument with SPRQ-Nx chemistry. ONT is from the Oxford Nanopore open-data GIAB 2025.01 release, flow cell PAW70337, basecalled with the super-accuracy model. Download locations for all six datasets are given in Availability of data and materials.

We processed all datasets identically through the allele-calling module. Within each comparison, datasets were downsampled to the minimum coverage among them so differences reflect the platform rather than depth. Downsampling used samtools view -s (samtools v1.16.1) [39,40].

#### Directional analysis of allele lengths between replicates

To ask whether replicate-unique alleles could reflect random jitter or directional bias between runs, we compared per-read allele lengths between HiFi-R23Q3a and HiFi-R23Q3b at 15,000 loci sampled uniformly at random from the loci passing the coverage filter above (at least 10 reads in each replicate). One-sided Mann-Whitney U test were used to test whether HiFi-R23Q3b lengths tend to exceed HiFi-R23Q3a lengths in each locus. Per-locus p-values across loci were corrected with the Benjamini-Hochberg procedure at a false discovery rate of 0.05. For per-locus effect size, we normalized the U statistic to U/(n1 n2), where n1 and n2 are the read counts for the two replicates. This quantity estimates the probability that a randomly chosen HiFi-R23Q3b read in that locus carries a longer allele than a randomly chosen HiFi-R23Q3a read. To test for a global directional shift, we compared the per-locus normalized U values against 0.5 using a one-sample, one-sided Wilcoxon signed-rank test, excluding loci with a value of exactly 0.5. Statistical tests were performed with SciPy (v1.15.3) [41], and multiple-testing correction used statsmodels (v0.14.5) [42].

#### Deriving error filters from technical replicates

For each ordered pair between HiFi-R23Q3a and HiFi-R23Q3b, and among HiFi-R23Q3, HiFi-S24 and HiFi-N26, we characterized every allele present in the test dataset but absent from the other by three measures: number of supporting reads, length offset from the closest allele at the same locus, and difference in motif k-mer count from that allele (Fig. 3e and Additional file 1: Figs. S5, S6, S7, S8 and S9). A mutation is reported only when the allele has at least 7 supporting reads within 9 bp of its length, is offset from the nearest allele in the comparison sample by at least 8 bp, and differs from it by more than 3 motif k-mers.

For large length changes, we did not filter by read support and instead required an offset of at least 100 bp from the nearest allele in the comparison sample. The requirement of more than 3 motif k-mers was retained. In the technical replicate comparison, alleles at offsets of 100 bp or more were sparse and were nearly all supported by a single read (Additional file 1: Fig. S16).

### Comparison with published de novo calls

#### Published call set

We took the de novo tandem repeat calls of Porubsky et al. [31] from their Supplementary Table 10 (sheet “TRs (GRCh38)”): 640 loci in the eight third-generation members, called with TRGT-denovo [30] and refined by Element short-read validation and manual review. Five of the eight members (NA12879, NA12881, NA12882, NA12885 and NA12886) have HiFi data in the pedigree release and were compared, covering 421 loci.

#### Rerunning TRumut on the published catalog

To remove catalog differences, we converted the platinumTRs v1.0 catalog used by Porubsky et al. (7,722,716 loci) to TRumut’s catalog format (chr, tr_start, tr_end, tr_motif) and annotated it with flanking uniqmers using the deposited uniqmer files [37]:

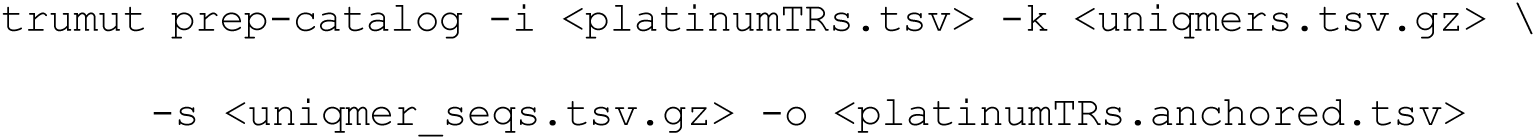

We then reran the allele-calling and trio modules on all 22 members with the parameters used elsewhere.

#### Matching calls

A TRumut de novo (clonal) call and a published locus were matched when they came from the same member and the anchored region overlapped the published interval. Several clonal rows at one region count as one locus, and a published locus listed with two de novo alleles counts once. No region overlapped more than one published locus.

#### Evaluating unmatched published loci

For each published locus without a TRumut call, we applied the trio module’s criteria to the per-read allele tables of the child and its parents (see “TRumut trio”). Each locus was classed by the first criterion it failed. A locus was classified as unassessable when a trio member had fewer than 10 reads, no read spanned both anchors, or the locus had no uniqmer on one side. Published size changes were taken from the child_to_parent_diff column of Supplementary Table 10 of Porubsky et al. Unassessable loci were intersected with the UCSC GRCh38 segmental duplication and centromere tables, with PAR1 taken as chrX:10,001-2,781,479, and the distance to the nearest uniqmer taken from the annotated catalog.

#### Measuring runtime and memory

We timed both pipelines on all 22 sequenced members of the CEPH 1463 pedigree and the 16 trios. The Tandem Repeats Finder catalog annotated with uniqmer anchors was used (see “Constructing the tandem repeat catalog”). All jobs ran on identical 64-core nodes of a Slurm cluster, one 8-CPU job per genome or trio. We ran TRGT v5.1.0 [1] in genotype mode with default parameters and the karyotype of each member. TRGT-denovo v0.3.0 [30] requires a sorted, indexed VCF and spanning-read BAM, which we produced with bcftools v1.19 and samtools v1.16.1 [39,40] before running it in trio mode:

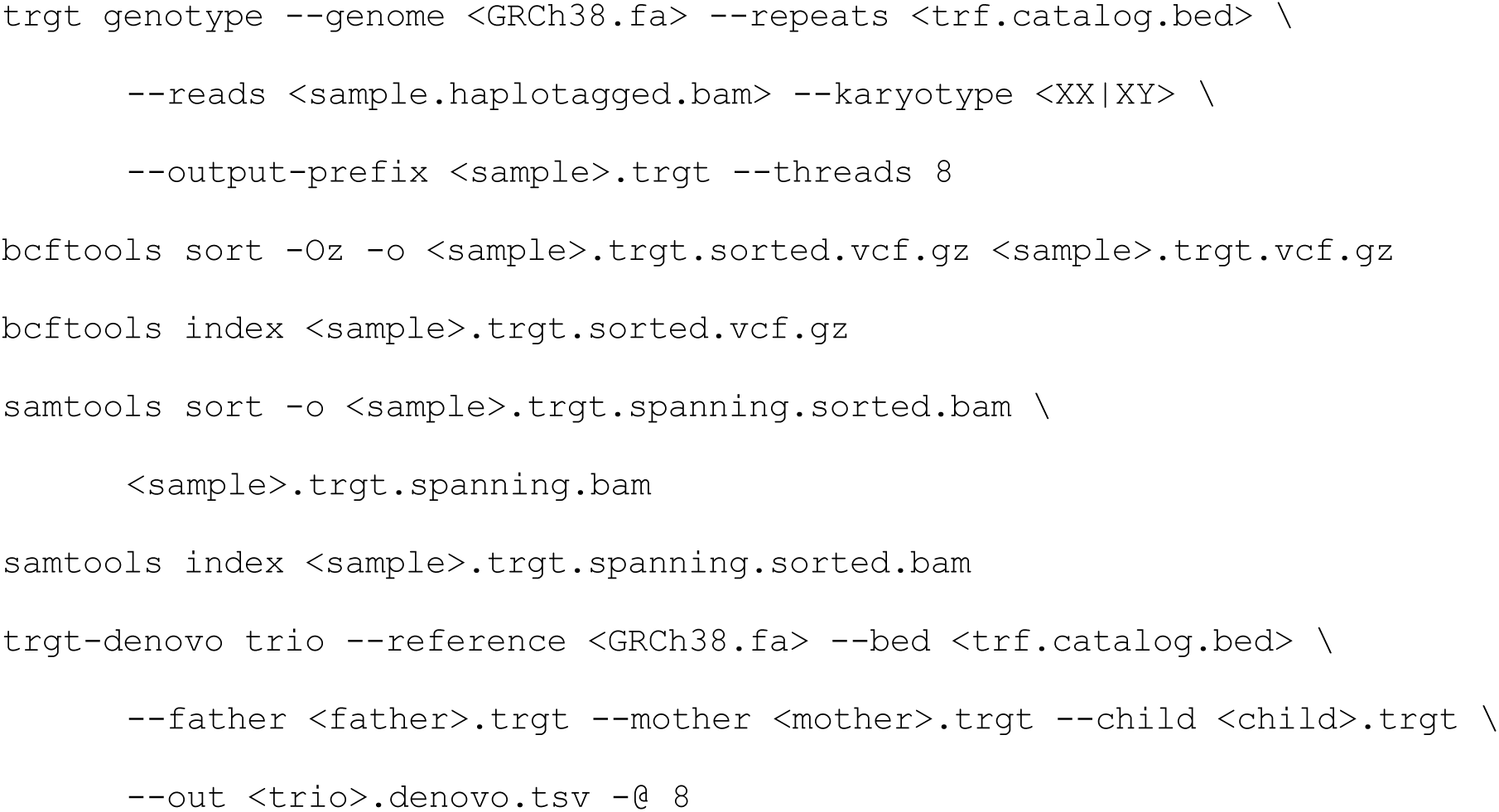

The TRumut pipeline comprised `trumut allele` and `trumut trio` with 8 workers each. We recorded CPU time and wall-clock time with GNU time for each command, and peak memory as the maximum resident set size summed over all processes of a job from Slurm accounting. We compared the two pipelines at three levels. Per genome, `trumut allele` was compared with TRGT genotyping including both sorts. Per trio, `trumut trio` was compared with trgt-denovo. End to end, each trio’s total was the per-genome time of its three members plus the trio step. We report CPU time because wall-clock time varied with co-scheduled jobs.

### TRumut trio

#### Calling nonparental alleles

The trio module takes the per-read allele tables of a child and its parents and reports child alleles absent from all of them. There is no limit in the number of parent samples, so that users can add relatives such as grandparents to constrain the inherited set further. At each anchored locus, allele lengths from all parental inputs are pooled. A child length is a candidate when it differs from the pooled parent length by at least the minimum offset (see “Deriving error filters from technical replicates”). Candidate lengths are then clustered greedily as in the `trumut mutual` module (see “Symmetric comparison with TRumut mutual”) with a wider merge width (7 bp). We analyze a locus only when the child and every parent input covering it each supply at least 10 reads. Each member was called against its parents with `trumut trio <child.alleles.tsv> <father.alleles.tsv> <mother.alleles.tsv> <child.trio.tsv> --input-type reads`, and large length changes by adding `--mega-events`, which sets the minimum offset to 100 bp and removes the read-support minimum.

#### Annotating and filtering candidates

Each candidate carries its signed offset (the distance to the nearest parent length), its supporting-read counts (reads within the support window of the length that also clear the minimum offset against the parents), and its median motif k-mer count, taken across the child reads at that length and compared with the median across parent reads at the nearest parent length (see “Motif k-mer counting”). TRumut then reports only the candidates passing the remaining two error-derived filters, both parameters defaulting to the values established from technical replicates (see “Deriving error filters from technical replicates”).

#### Classifying event types

The trio module labels each event clonal, subclonal, or private, as defined in Results. We report clonal alleles as de novo and subclonal and private alleles as postzygotic.

For each event, the module gathers the child reads whose lengths fall within the read-support window. The haplotype tag carried by most of those reads becomes the allele’s haplotype, and untagged reads count toward it when the window contains no reads of the other haplotype. The phased classification is used only when at least half of the child’s reads at the locus carry a haplotype tag and at least two tagged reads fall within the window. Otherwise, the allele takes the unphased path described below.

An allele supported by a single read is automatically classified as private. Otherwise, the module examines every read of that haplotype within the read-support window of the reported length, some of which has lengths that do not clear the minimum offset against the parents. Of those that do, the one with the most reads represents the allele. The module then compares that length with every other length in the window of that representative, including those too close to a parental length to count as deviating. If one has more reads, the allele cannot be separated and is classified as private.

The remaining alleles are separated into clonal and subclonal. A de novo allele should appear on every read of its haplotype, so a haplotype carrying a second distinct allele indicates a postzygotic origin. We use two tests to check if a second allele exists on that haplotype, and passing either one marks the allele as subclonal. First, lengths that lie within the read-support window of each other are chained, so that one allele’s jitter stays together. We then ask whether a chain that does not contain the representative holds at least four reads or one tenth of the haplotype’s reads, whichever is larger. Second, the haplotype’s lengths are smoothed with the same window. They are then evaluated whether the densest point lies more than a window away from the representative with that much support, or whether another supported length more than a window away is separated from the representative by a dip to half or less of the smaller of the two peaks. The first test is meant to catch well-separated alleles, and the second a minority allele bridged to the main one by a sparse tail. An allele that fails both is clonal.

When the window holds no haplotagged read or the locus fails the tagging requirements above, the same tests are run but with all reads pooled. We then label the allele only as private or “not private, unresolved.” Without a haplotype, the module cannot ask whether a second allele exists, and therefore cannot distinguish clonal from subclonal events. The private test is also less reliable without phase, because reads from both haplotypes compete in the same comparison.

#### Trio data

We applied the trio module across the CEPH 1463 platinum pedigree, using HiFi data from 22 sequenced members [31]. Each member with sequenced parents was analyzed against them. Reads were processed with the allele-calling module as above. Haplotype tags were used as released, assigned with WhatsHap v1.4 [43] against a phased DeepVariant call set [44].

#### Transmission of de novo alleles

For each de novo event in a member with sequenced children, we identified the allele each child received from the carrier parent using the released phasing. Each child haplotype was assigned to whichever parent carried the allele closer to it in length. The haplotype assigned to the carrier was then classified as the de novo allele, the carrier’s other allele, or neither. In the process, two allele lengths must differ by less than the minimum offset to be determined as matching.

#### Identifying recurrent loci

To ask whether large length changes recur at particular loci, we intersected events across members. A locus was considered recurrent when mutations were called in two or more members spanning at least two distinct trios. We then classified recurrent loci by pedigree structure. A locus was defined as “intra-generationally recurrent” when two or more members of the same generation carried an event, and “inter-generationally recurrent” when at least one pair of carriers belonged both to different generations and to different trios. These categories are not mutually exclusive. We only performed recurrence analyses on the 16 third- and fourth-generation members whose sequencing depths are most similar.

For each recurrent locus, we assessed concordance in two ways. First, we tested whether the events agreed in direction, all being expansions or all contractions. We computed the difference in allele length between the mutated allele’s representative read and the closest parental read, and took its sign as the direction. Events where this difference was within 7 bp were left out of the comparison. Second, we asked whether the mutated alleles also matched in length. A locus matched in length when every pair of mutated allele lengths from different members agreed within 7 bp. It must also match in direction. For intra-generational recurrence, we assessed concordance within each generation separately.

### TRumut cancer

The cancer module calls somatic tandem repeat alleles in a tumor against its matched normal. It shares the trio module’s machinery, with the tumor as the test and the matched normal as comparison. Tumor lengths that deviate from every normal length by at least the minimum offset are clustered into candidates, which are filtered and annotated as described above. Each reported allele is classified as clonal, subclonal, or private using the same read-support and phasing rules. We use the term “clonal” to refer to truncal mutations, “subclonal” for tumor subclones, and “private” for those that are too sparse to constitute an independent allele cluster. Because structural variant breakpoints often fall in tandem repeats, non-motif k-mer localization (see “Localizing non-motif indels relative to the repeat”) is particularly useful here. Each tumor was called with `trumut cancer <tumor.alleles.tsv> <normal.alleles.tsv> <tumor.cancer.tsv> --input-type reads`, and large length changes with `--mega-events` as for the trio module.

#### Cancer data

We applied the TRumut cancer module to six HiFi tumor-normal pairs (COLO829, HCC1395, HCC1937, HCC1954, H1437, and H2009), aligned to GRCh38. We called small variants in every tumor and normal separately with DeepVariant v1.6.1 [44] and haplotagged reads with HiPhase v1.7.0 [45], requiring a minimum variant quality of 20 and disabling global realignment and read-group handling:

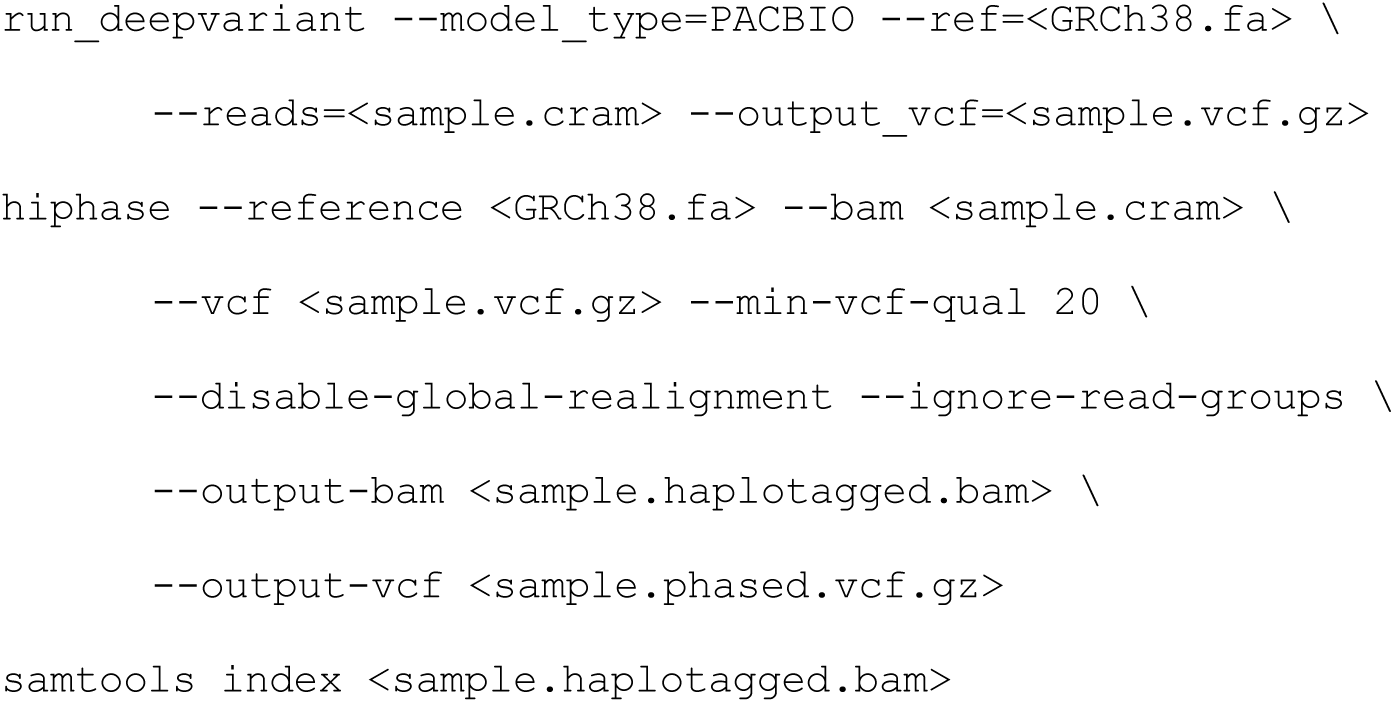

### TRumut use-pon

The `trumut use-pon` module calls extreme alleles in a sample with no matched normal or sequenced parents. Instead, it uses the allele lengths carried by the population at each locus as the comparison. We applied the `trumut use-pon` module to 10 cancer cell lines, the six tumors above and four lines without matched normals (see Availability of data and materials).

#### Building a panel of normals

We built the panel from the reads of 232 HPRC genomes, each processed with the allele-calling module and the same catalog. The 232 per-sample tables were merged into one entry per anchored locus with `trumut build-pon <alleles_dir/> <pon.tsv> --input-type reads`. A genome “covers” a locus when it supplies at least 10 distinct reads there, and we include a locus in the panel only when at least 98% of genomes cover it. This requirement dropped 0.6% of loci.

For each locus that enters the panel, we record the range of allele lengths supported among genomes that cover that locus. A length carried by fewer than 9.75% of the covering genomes is excluded from the range. This cutoff is calculated to represent allele frequencies of 5% under Hardy-Weinberg equilibrium. Rare variants are thus excluded since they would widen the range and mask test-sample alleles of the same length. At each end of the range, we also record the median motif k-mer count of reads carrying that length, so that motif content can be compared although the panel retains no reads.

#### Calling extreme alleles against the panel

The allele length of every read from the test sample is compared against the “normal” length range recorded in the panel. A read is kept only when its length falls at least 8 bp beyond either end of the panel range.

Kept lengths are clustered as in the mutual module (see “Symmetric comparison with TRumut mutual”) and held to the same error-derived thresholds as the other callers (see “Deriving error filters from technical replicates”), with motif k-mer content compared against the panel’s recorded median at the nearer end of the range (see “Building a panel of normals”).

Samples were called against the panel with trumut use-pon <sample.alleles.tsv> <pon.tsv> <sample.pon.tsv> --input-type reads.

#### Outlier allele detection

The per-read tables of the 10 lines were screened for alleles far from every other allele at the locus, without a matched normal or panel. At each anchored locus, we sorted the per-read allele lengths reported by the allele-calling module. An allele was called an “outlier” if it is the longest at the locus with the second-longest at least 100 bp shorter, or if the shortest with the second-shortest at least 100 bp longer. Outliers were counted per line and normalized by coverage. HG002, processed identically, is the negative control.

#### Comparison with microsatellite instability status

We obtained microsatellite instability scores from the DepMap portal. We used the MSI score field in OmicsGlobalSignatures.csv and mapped model identifiers to cell line names using Model.csv. When a cell line had multiple entries, we took the average MSI score.

## Declarations

### Ethics approval and consent to participate

Not applicable.

### Consent for publication

Not applicable.

### Availability of data and materials

TRumut is openly available at https://github.com/erkim11/TRumut. The uniqmer sets, the anchored tandem repeat catalog, and the panel of normals are available at Zenodo, https://doi.org/10.5281/zenodo.22903449 [37]. Mutation calls underlying all figures are available at Zenodo, https://doi.org/10.5281/zenodo.23444518 [46]. Per-read allele tables can be regenerated from the datasets listed below using TRumut and the deposited catalog.

HiFi sequencing data for the four cancer cell lines that did not have matched normals are deposited in the NCBI Sequence Read Archive under BioProject PRJNA1554400, https://www.ncbi.nlm.nih.gov/sra/PRJNA1554400.

All other datasets analyzed in this study are publicly available. HG002: HiFi-R23Q1, https://human-pangenomics.s3.amazonaws.com/submissions/80d00e88-7a92-46d8-88c7-48f1486e11ed--HG002_PACBIO_REVIO/m84039_230117_233243_s1.hifi_reads.default.bam [47], HiFi-R23Q3a and HiFi-R23Q3b, https://ftp-trace.ncbi.nlm.nih.gov/ReferenceSamples/giab/data/AshkenazimTrio/HG002_NA24385_son/PacBio_HiFi-Revio_20231031/ [48], HiFi-S24, https://downloads.pacbcloud.com/public/revio/2024Q4/WGS/GIAB_trio/HG002_rep1/ [49], HiFi-N26, https://downloads.pacbcloud.com/public/2026Q2/HG002-SPRQ-Nx/ [50], and ONT, https://epi2me.nanoporetech.com/giab-2025.01/ [51]. CEPH 1463 pedigree HiFi data, https://registry.opendata.aws/platinum-pedigree/ [31]. HiFi data for the six tumor-normal pairs were obtained from the NCBI Sequence Read Archive under BioProject PRJNA1086849 (H1437, H2009, HCC1937, HCC1954) [52] and from PacBio public datasets (COLO829, https://downloads.pacbcloud.com/public/revio/2024Q4/WGS/COLO829/; HCC1395, https://downloads.pacbcloud.com/public/revio/2023Q2/HCC1395/) [53,54]. Microsatellite instability scores, DepMap Public 26Q3, https://depmap.org/portal [55,56].

HiFi reads for the HPRC genomes were obtained from the HPRC data portal, https://data.humanpangenome.org/raw-sequencing-data [57]. Haplotype-resolved assemblies for uniqmer selection were obtained from the Open Human Genome Library v4.0, https://registry.opendata.aws/openhgl/, archived at https://doi.org/10.5281/zenodo.13948741 [29], which collects assemblies from HPRC Release 2 [32], the Arab Pangenome Reference [58], CN1 [59], KSA001 [60], I002C [61], KOREF1 [62], YAO [63], and CHM13 [64,65].

### Authors’ contributions

ERK and HL designed the method and the analyses. ERK implemented the method and wrote the manuscript. HL supervised the project and revised the manuscript. Both authors read and approved the final manuscript.

## Acknowledgements

We thank Jim Shaw, Ying Zhou, Jakob Heinz, Megan Le, and other members of the Li lab for helpful discussions.

## Supplementary Information

Additional file 1: Figures S1-S18.

Additional file 2: Table S1. Transmission of de novo alleles to sequenced children in the CEPH 1463 pedigree.

Additional file 3: Table S2. Published de novo TR loci at which TRumut withheld a call.

