## Additional file 1: Figures S1-S18 for "Pangenome-conserved k-mers enable accurate de novo and somatic tandem repeat mutation calling from long reads"

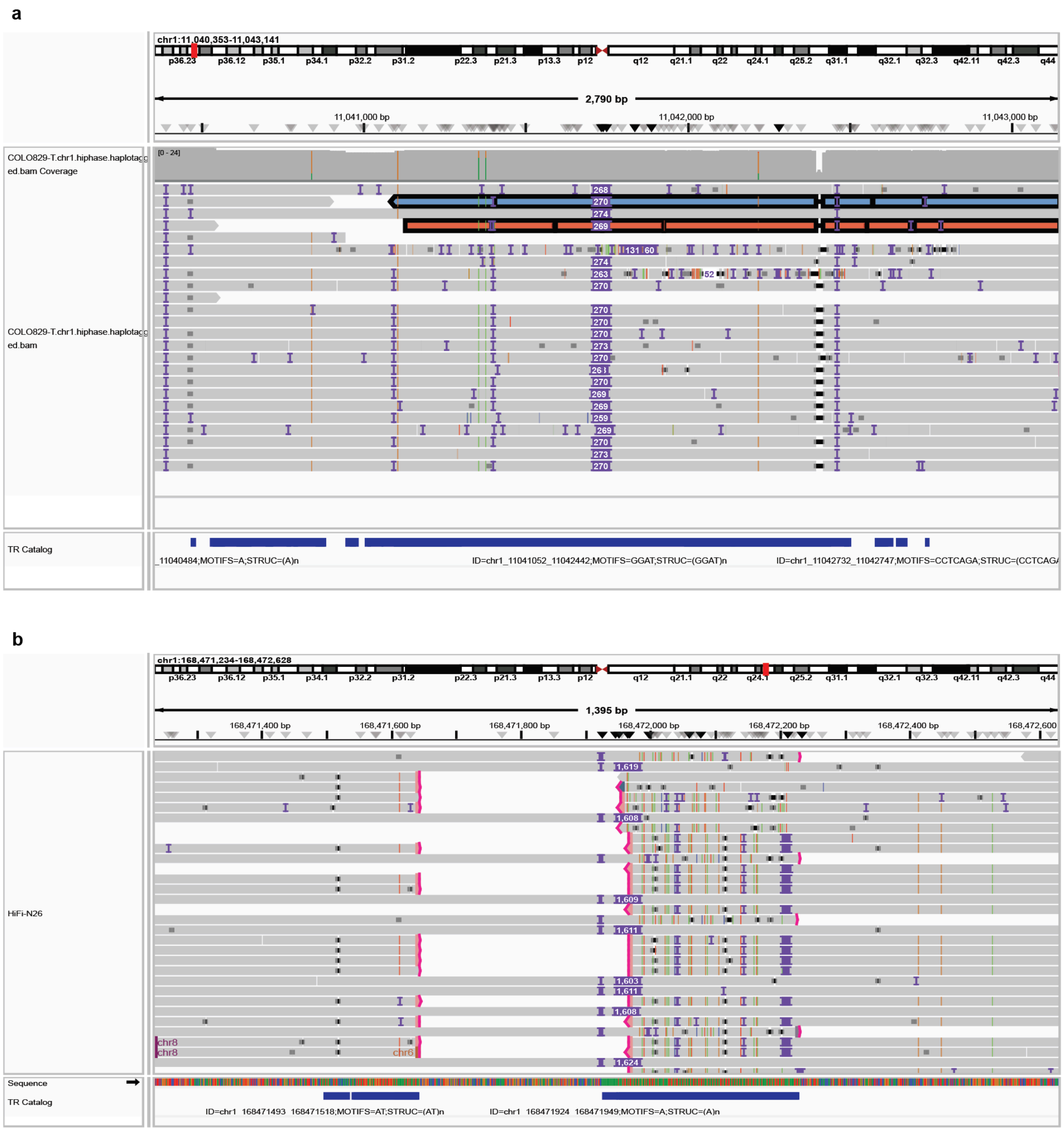


**Figure S1. Alignment failures at tandem repeat loci in HiFi reads.**

IGV views of HiFi alignments. Purple boxes give insertion lengths in bp; pink marks soft-clipped read ends.

a. chr1:11,040,353-11,043,141 in COLO829 tumor. Most reads carry a ~270 bp insertion in the GGAT repeat at chr1:11,041,052-11,042,442. The two highlighted reads begin inside the repeat and do not reach the left flank, yet sequence within the repeat matches the flank at ≥70% identity and is aligned as if it were the flank, so the allele is determined to be far shorter than the repeat (Fig. 1b, Underestimated).

b. chr1:168,471,234-168,472,628 in HiFi-N26. Every read carries a ~1.6 kb insertion at the A homopolymer at chr1:168,471,924-168,471,949. Some reads represent it as a single insertion of 1,603-1,624 bp; the rest are soft-clipped on both sides of it, with the clipped segments mapped to other chromosomes (chr6, chr8). The clipped reads have no aligned flank on one side, so no allele is called from them (Fig. 1b, No call), and the two representations assign different lengths to the same allele.

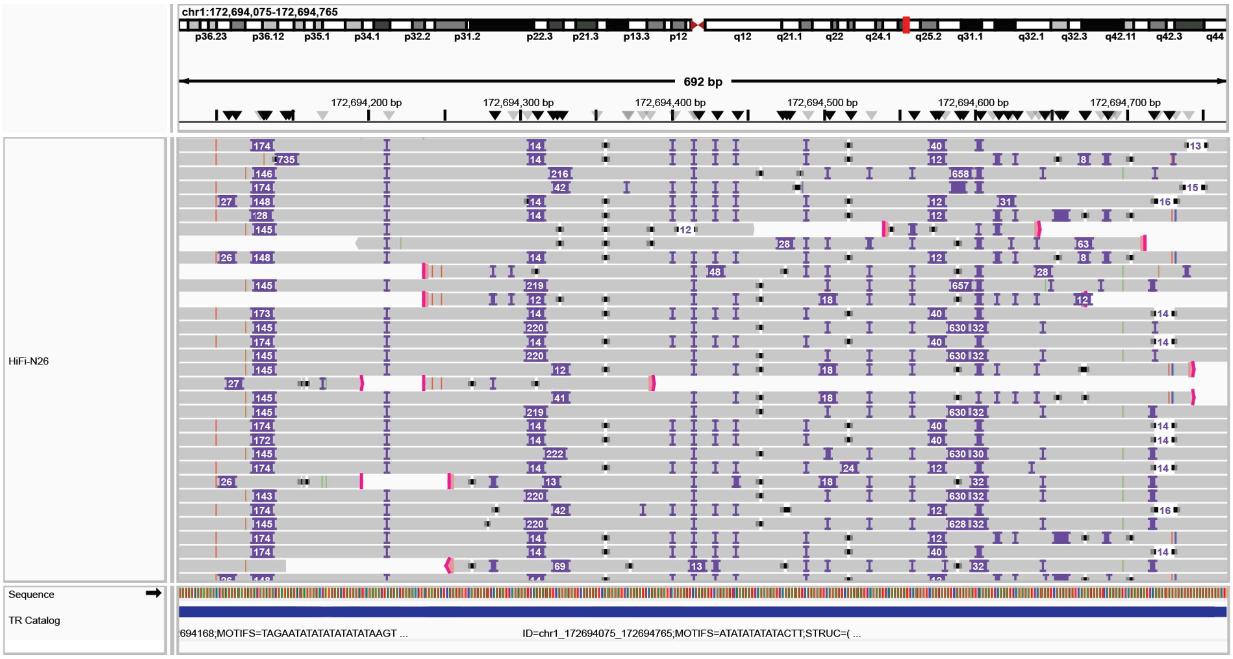


**Figure S2. Inconsistent alignment within a tandem repeat.**

chr1:172,694,075-172,694,765 in HiFi-N26, an AT-rich VNTR spanning the catalog interval. IGV notation as in Fig. S1. Reads from the same haplotype are aligned very differently from one another. Some reads are clipped mid-repeat.

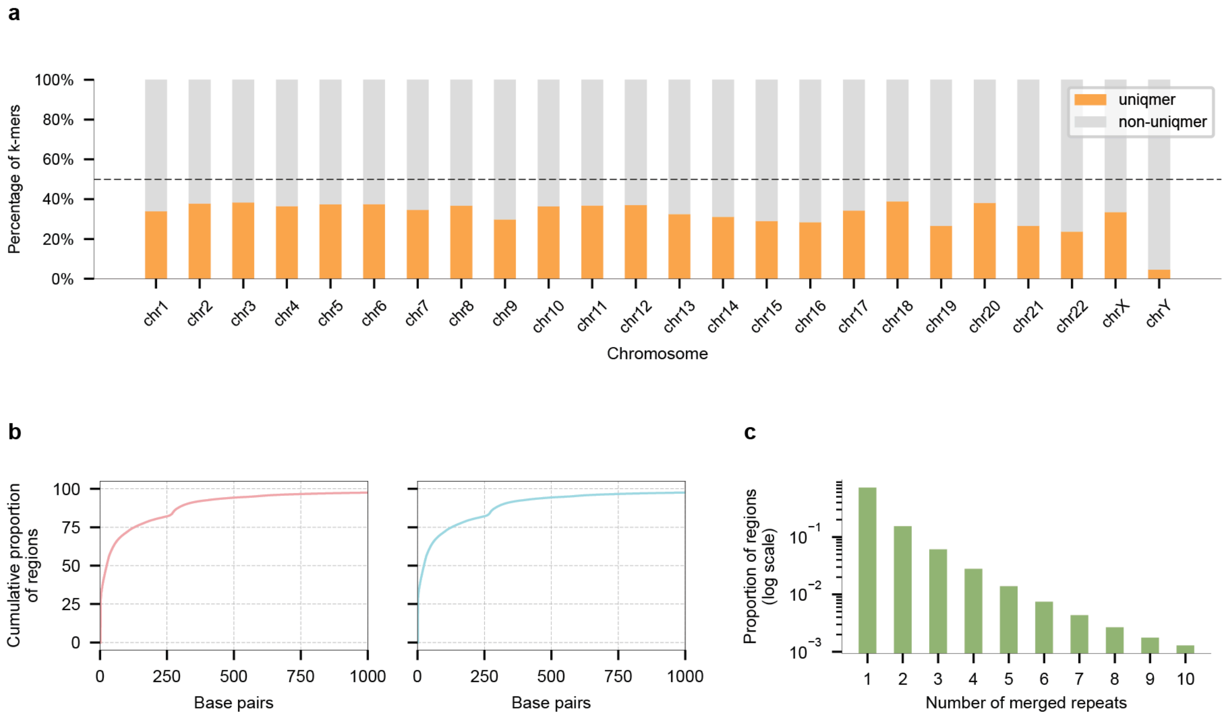


**Figure S3. Properties of the uniqmer-anchored catalog.**

a. Proportion of all 31-mers that qualify as uniqmers, per chromosome. Centromeric and telomeric regions were excluded.

b. Cumulative proportion of anchored regions by the distance from the repeat to its closest uniqmer, measured at the start (left) and end (right) of the repeat. Distances are in base pairs, truncated at 1 kb.

c. Number of repeats merged into a single anchored region, on a log-scaled frequency axis. Most regions required no merging.

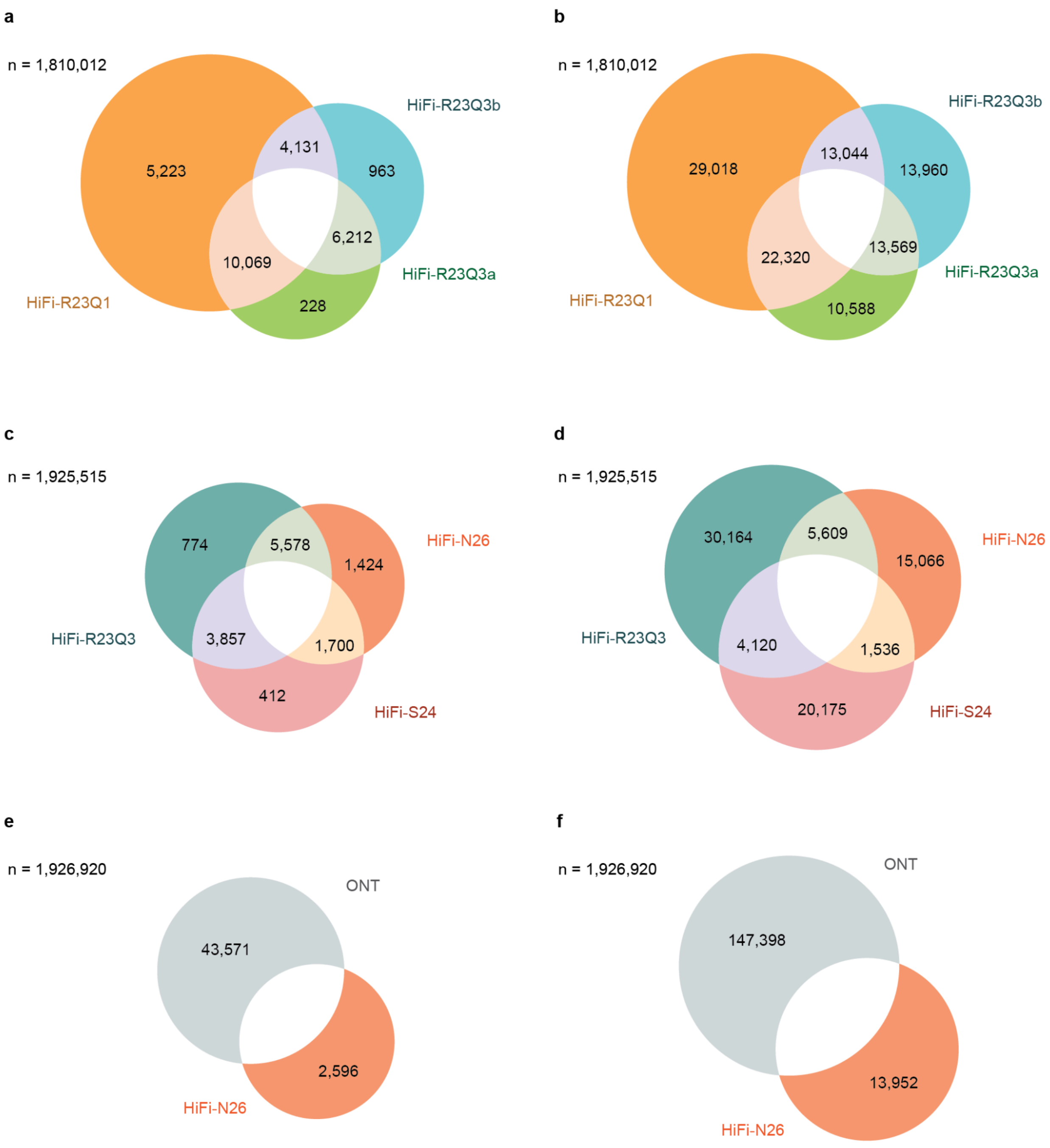


**Figure S4. Replicate comparisons under stricter criteria.**

Loci at which at least one allele was detected in one dataset but absent from another, as in Fig. 3a-c. Left column (a, c, e), counting only alleles supported by at least two reads of exactly that length. Right column (b, d, f), counting only alleles more than 6 bp from any allele in the other datasets. a, b, HiFi-R23Q1, HiFi-R23Q3a, and HiFi-R23Q3b at 24x. c, d, HiFi-R23Q3, HiFi-S24, and HiFi-N26 at 40x. e, f, HiFi-N26 against ONT at 40x.

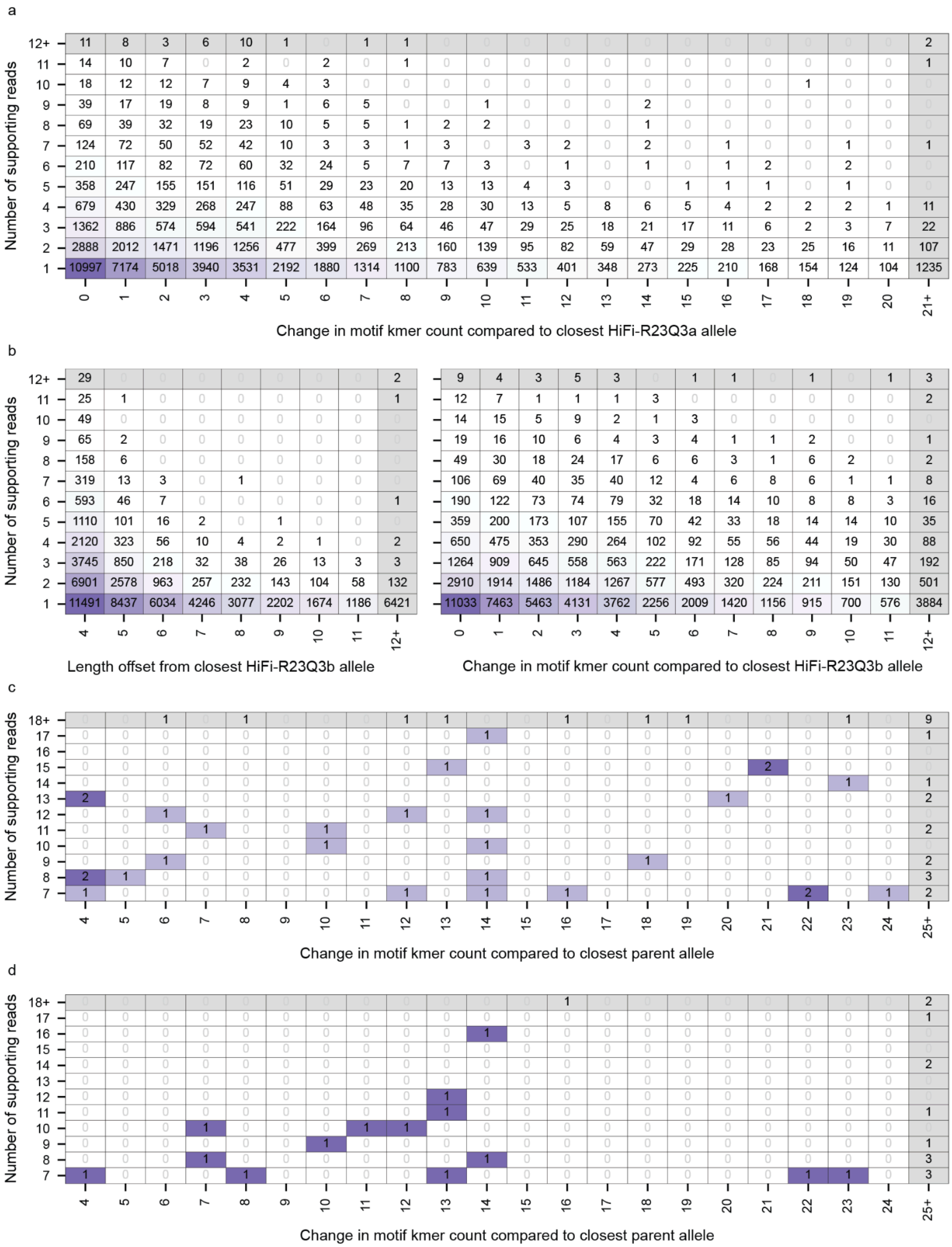
a
Number of supporting reads
Change in motif kmer count compared to closest HiFi-R23Q3a allele
b
Number of supporting reads
Length offset from closest HiFi-R23Q3b allele
Change in motif kmer count compared to closest HiFi-R23Q3b allele
c
Number of supporting reads
Change in motif kmer count compared to closest parent allele
d
Number of supporting reads
Change in motif kmer count compared to closest parent allele

**Figure S5. Replicate-unique and mutant alleles cross-tabulated by motif k-mer count, and the reciprocal replicate comparison.**

Layout and conventions as in Fig. 3e: cell values and shading give allele counts, and the final row and column aggregate larger values. Motif k-mer difference is the absolute difference between the allele's median motif k-mer count and that of the closest allele in the comparison sample (Methods).

a. Alleles in Fig. 3e, detected in HiFi-R23Q3b but absent from HiFi-R23Q3a, by supporting reads and motif k-mer difference.
b. The reciprocal comparison, alleles detected in HiFi-R23Q3a but absent from HiFi-R23Q3b, by supporting reads and, left, length offset or, right, motif k-mer difference.
c. De novo events in fourth-generation pedigree members (Fig. 4c), by supporting reads and motif k-mer difference.
d. Postzygotic events in fourth-generation pedigree members (Fig. 5c), by supporting reads and motif k-mer difference.

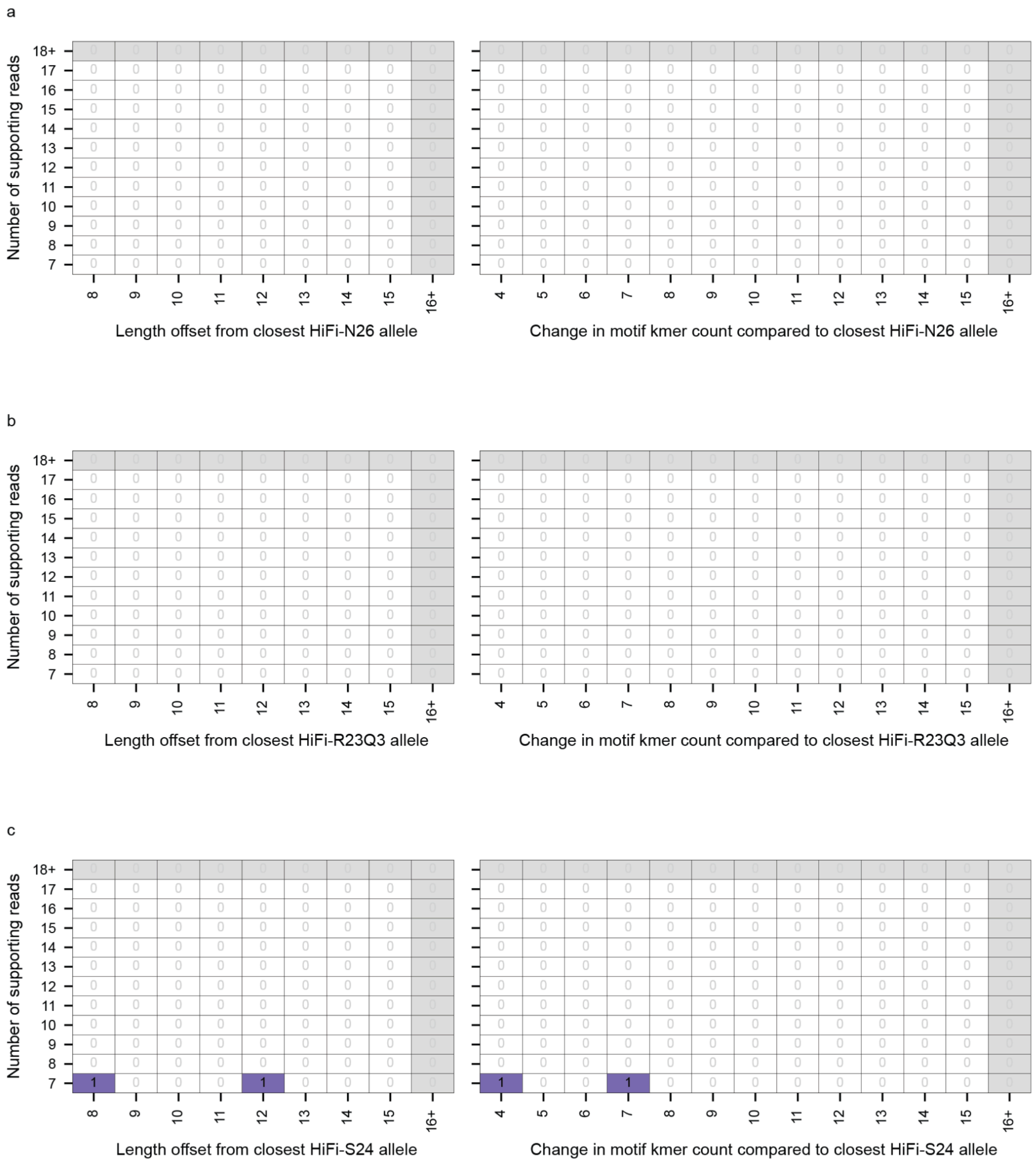


**Figure S6. De novo criteria applied between HiFi datasets of the same genome.**

Alleles present in one HiFi dataset but absent from a second, filtered with the thresholds used for de novo calls and retained only when classified as clonal (Methods). All datasets derive from HG002, so surviving alleles are more likely to reflect error than true de novo mutation. Each panel cross-tabulates the surviving alleles by the number of supporting reads and, left, the length offset from the closest allele in the comparison dataset or, right, the difference in motif k-mer count from that allele. Cell values and shading give allele counts, and the final row and column aggregate larger values.

a. Present in HiFi-R23Q3, absent from HiFi-N26.
b. Present in HiFi-S24, absent from HiFi-R23Q3.
c. Present in HiFi-N26, absent from HiFi-S24.

Reciprocal comparisons are shown in Fig. S7.

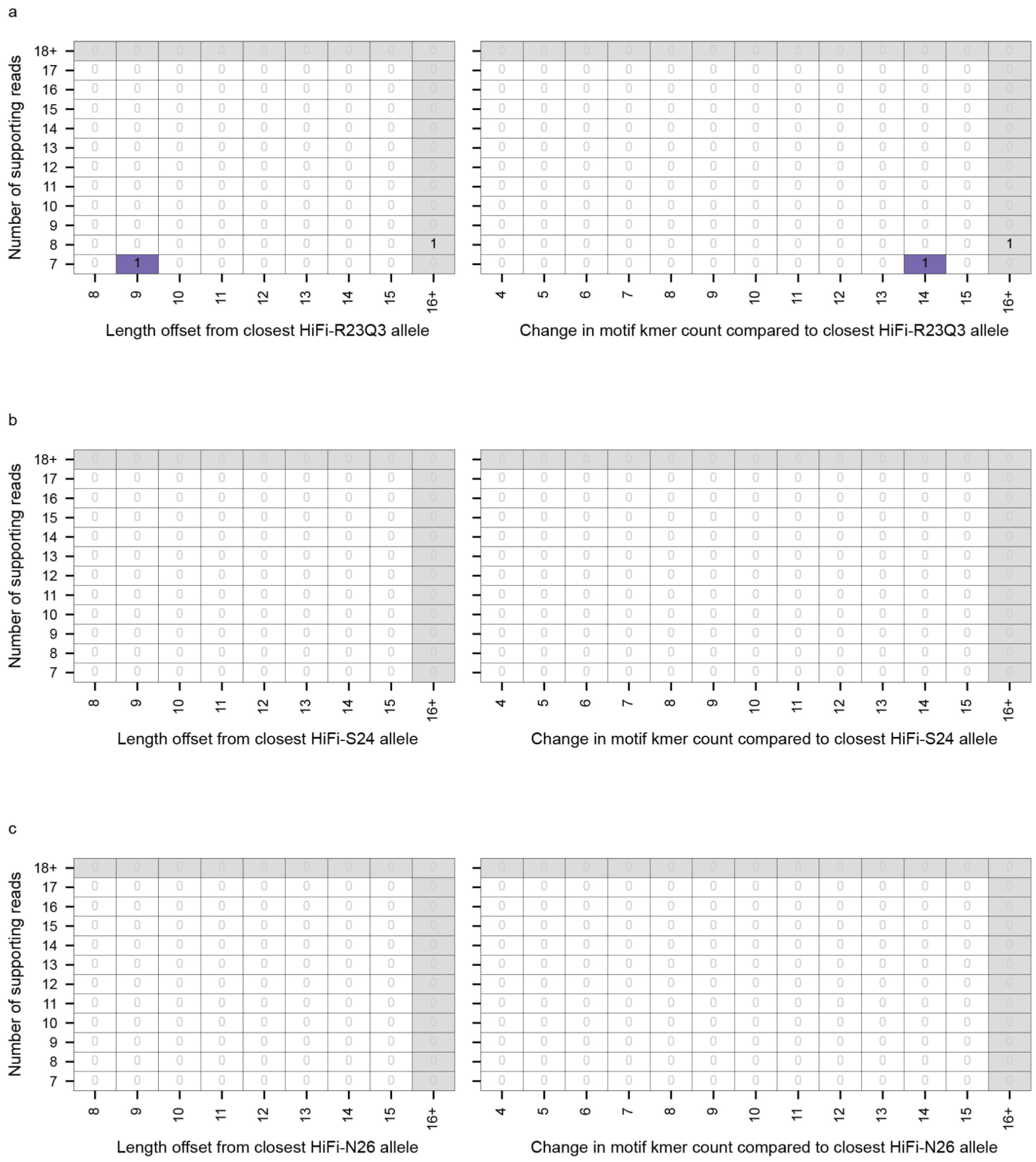


**Figure S7. De novo criteria applied between HiFi datasets of the same genome, reciprocal comparisons.**

As in Fig. S6, with the roles of the two datasets exchanged.

a. Present in HiFi-N26, absent from HiFi-R23Q3.

b. Present in HiFi-R23Q3, absent from HiFi-S24.

c. Present in HiFi-S24, absent from HiFi-N26.

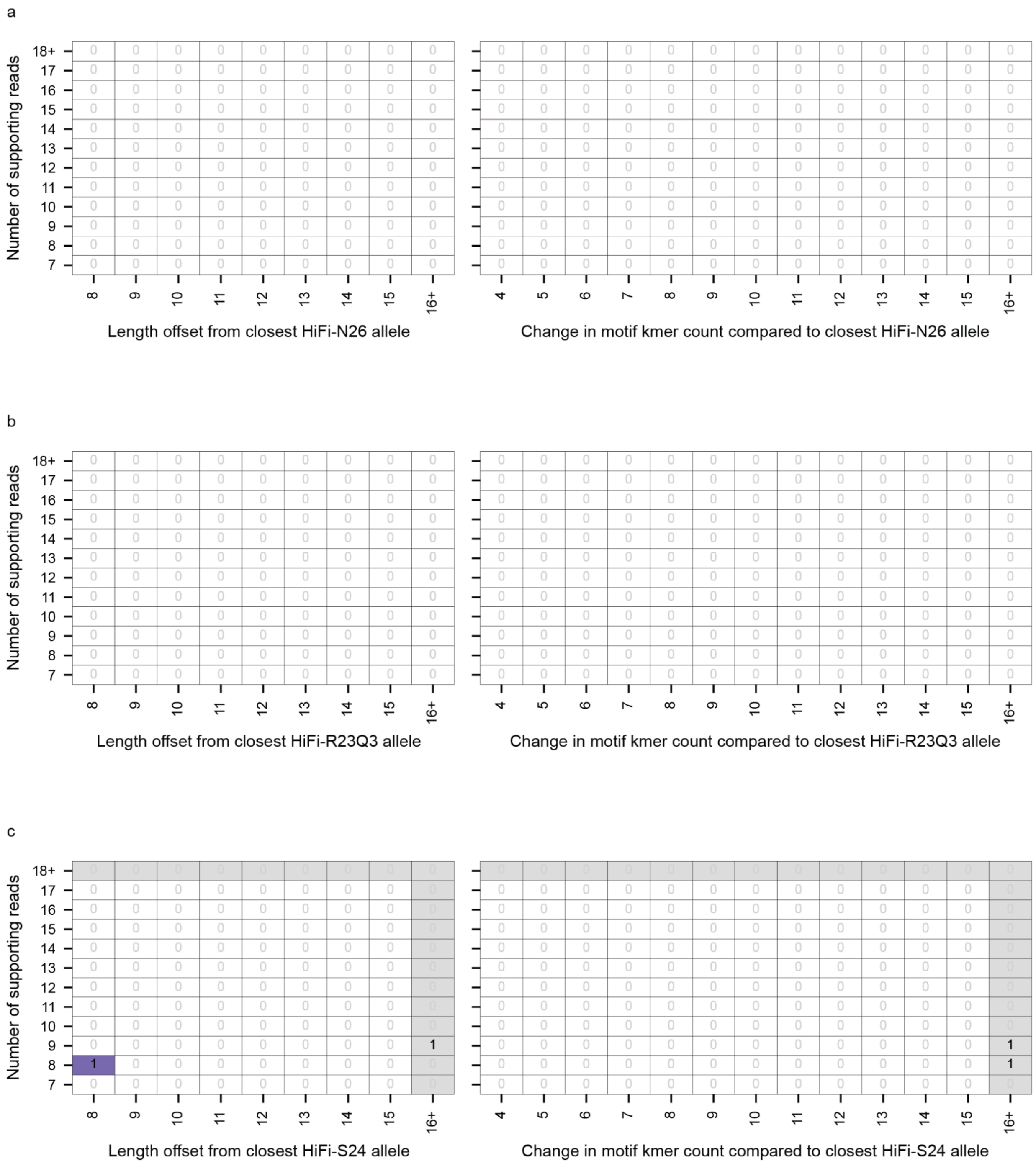

**Figure S8. Somatic criteria applied between HiFi datasets of the same genome.**

As in Fig. S6, but filtered with the thresholds and classes applied to postzygotic and cancer calls (Methods).

a. Present in HiFi-R23Q3, absent from HiFi-N26.

b. Present in HiFi-S24, absent from HiFi-R23Q3.

c. Present in HiFi-N26, absent from HiFi-S24.

Reciprocal comparisons are shown in Fig. S9.

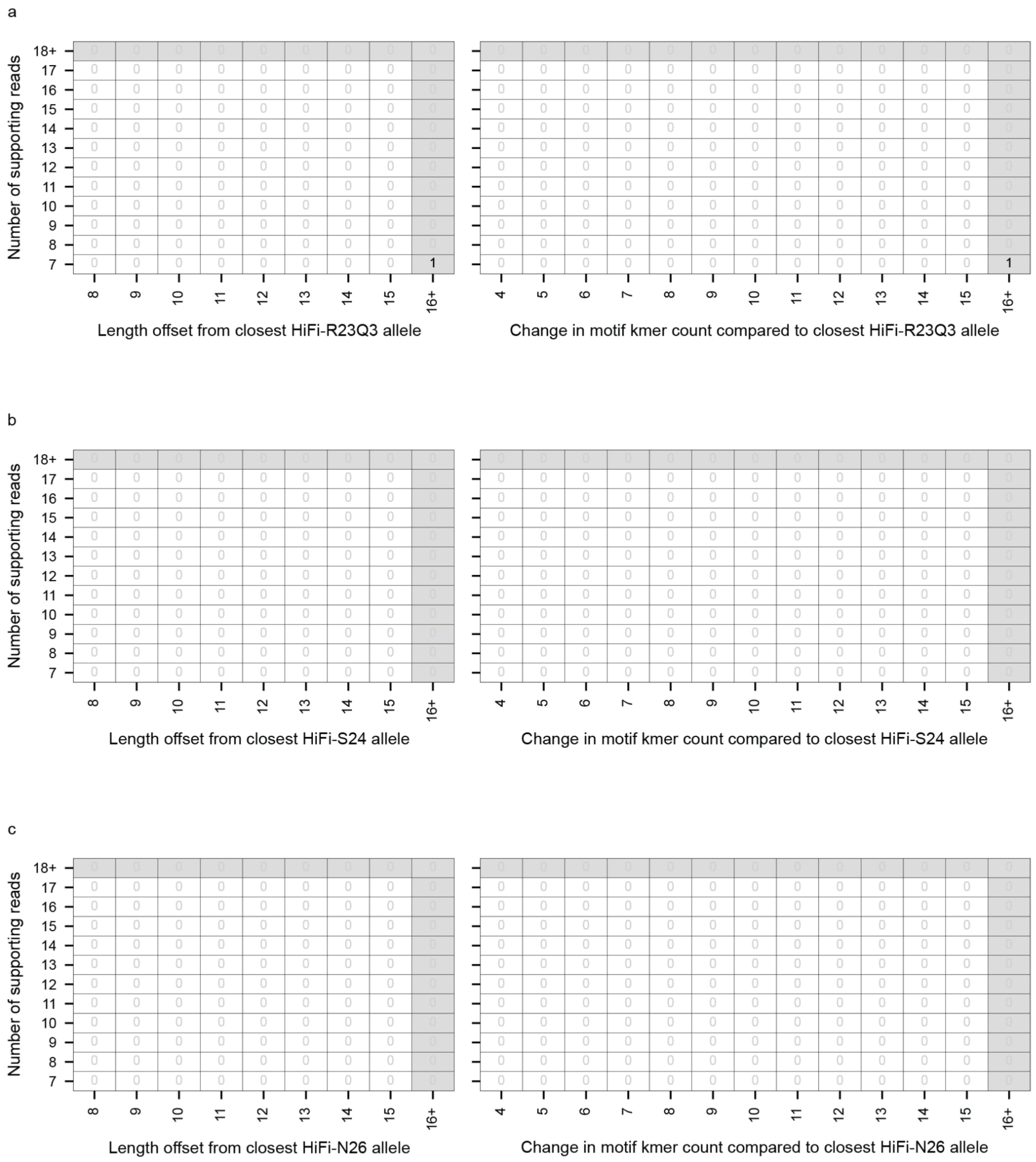


**Figure S9. Somatic criteria applied between HiFi datasets of the same genome, reciprocal comparisons.**

As in Fig. S8, with the roles of the two datasets exchanged.

a. Present in HiFi-N26, absent from HiFi-R23Q3.

b. Present in HiFi-R23Q3, absent from HiFi-S24.

c. Present in HiFi-S24, absent from HiFi-N26.

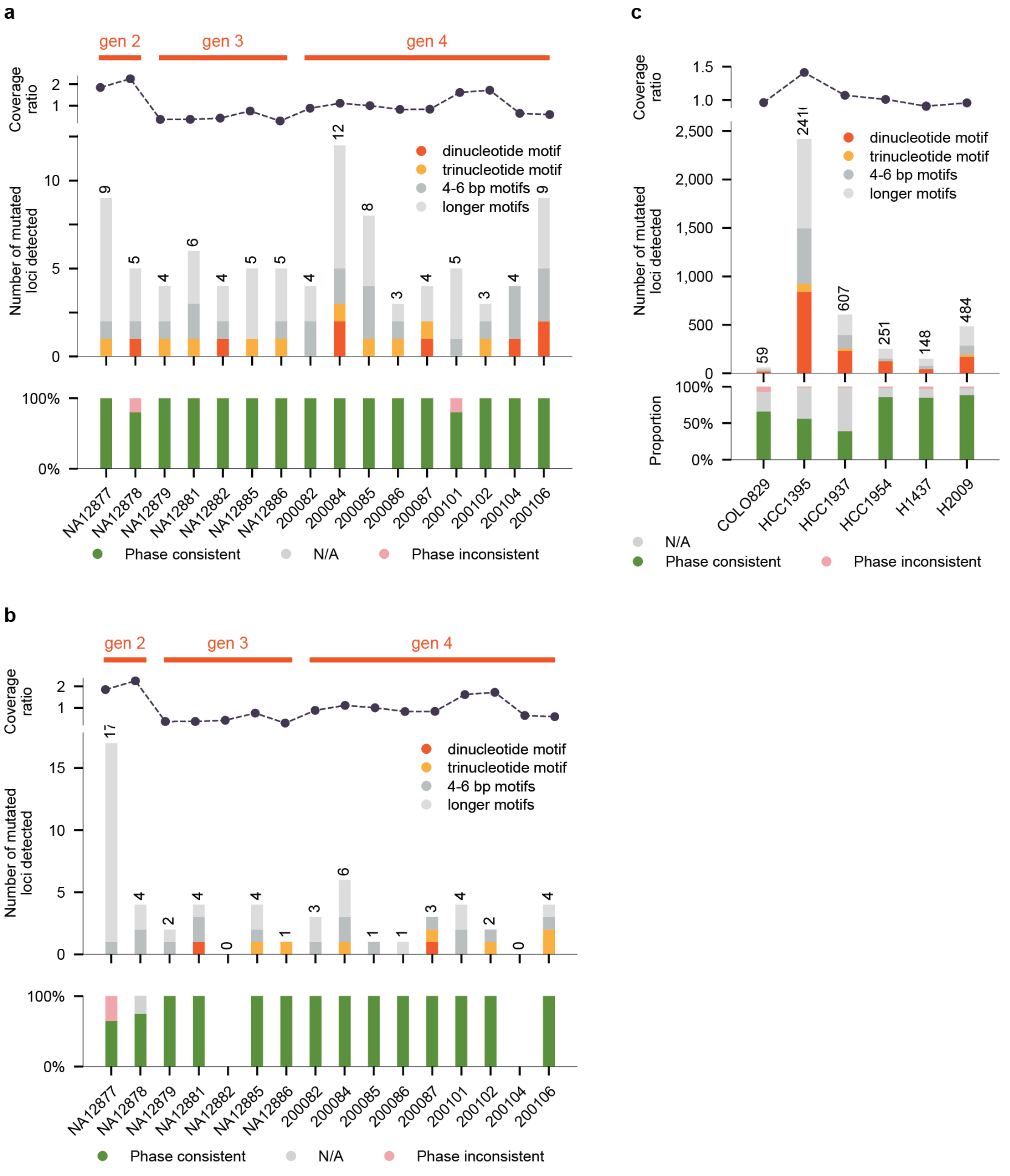


**Figure S10. Mutated loci by repeat motif length.**

As in Figs. 4b, 5b and 6b, with loci colored by the length of the repeat motif rather than by length offset. For merged regions, the motif spanning the most reference sequence is used (Methods).

Motif classes are dinucleotide (2 bp), trinucleotide (3 bp), 4-6 bp, and longer motifs (7 bp or more). Homopolymers were excluded from all analyses, so no single-base class appears. Only loci with an allele called in at least 10 reads in every dataset compared were analyzed (Methods).

a. As in Fig. 4b, de novo loci per pedigree member.
b. As in Fig. 5b, postzygotic loci per pedigree member.
c. As in Fig. 6b, somatic loci per cancer cell line.

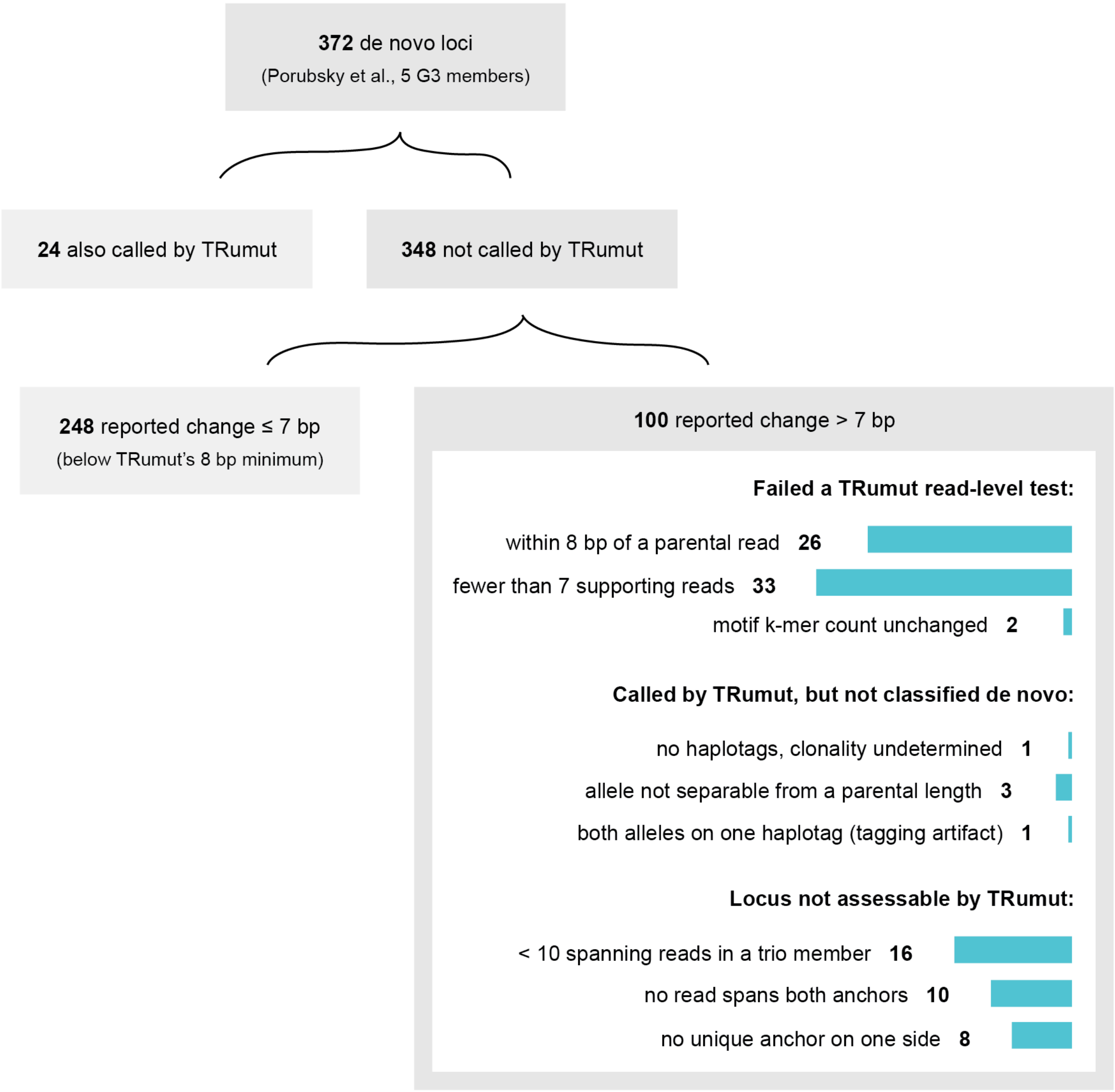


**Figure S11. Comparison with TRGT-denovo.**

Fate of the 372 de novo loci that Porubsky et al. reported with TRGT-denovo in the five third-generation members of the CEPH 1463 pedigree. TRumut was rerun on the same catalog and reported 24. Of the 348 it did not, 248 had a reported change of 7 bp or less, below our 8 bp offset threshold. The other 100 are grouped by reason.

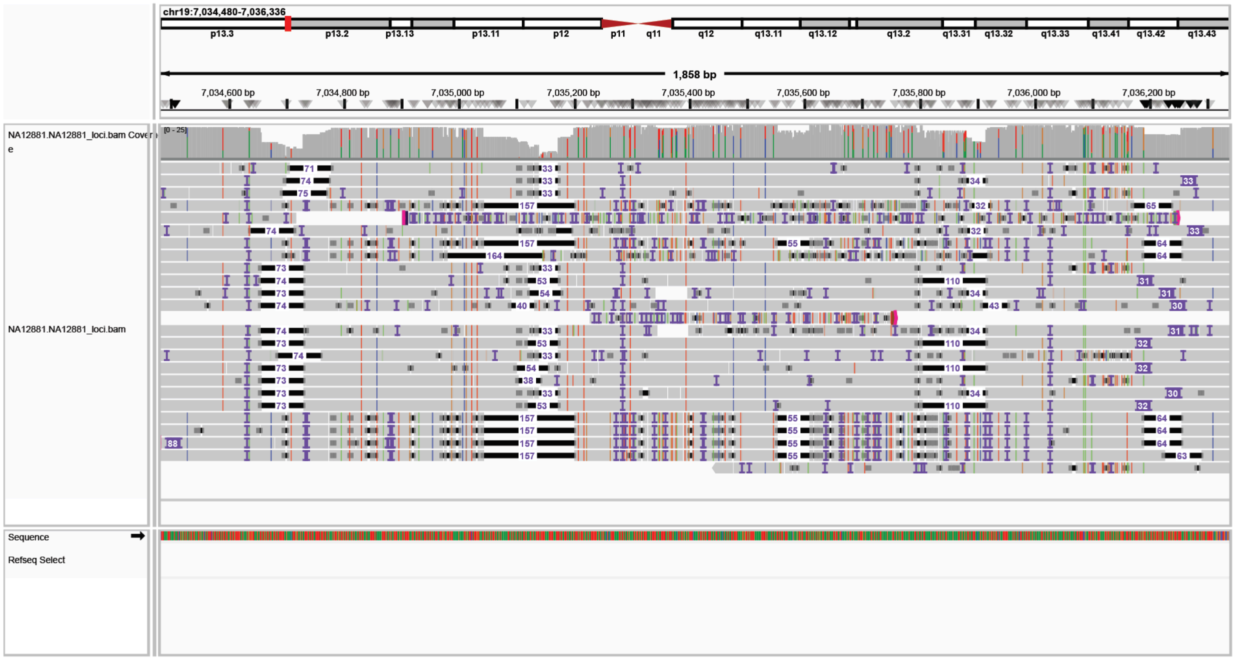


**Figure S12. Inconsistent alignment at a published de novo TR locus.**

chr19:7,034,449-7,036,305 in NA12881, an AT repeat inside a segmental duplication, reported by Porubsky et al. as a -91 bp paternal de novo allele. IGV notation as in Fig. S1. Reads map with MAPQ 60, yet reads from the same haplotype are aligned in markedly different ways. Two reads are clipped mid-repeat with up to 20 kb left unaligned.

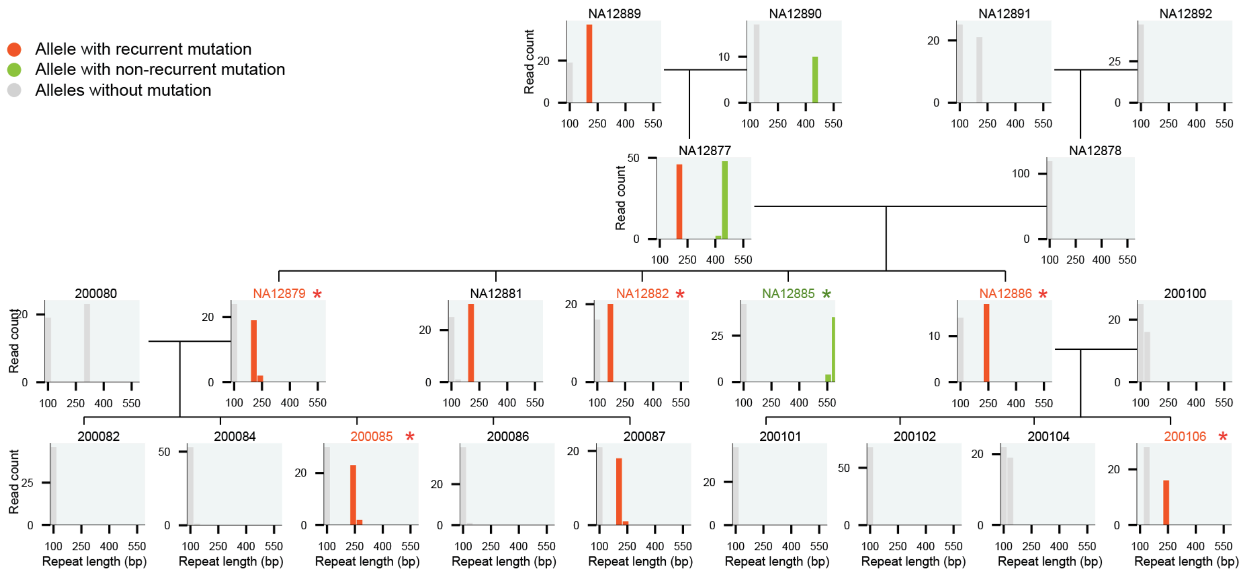


**Figure S13. A tandem repeat allele mutated in multiple generations.**

Read-level allele length distributions at chr8:2,623,352-2,623,432 for every sequenced member of the pedigree, arranged by pedigree position. Color marks one lineage tracked through the pedigree: red, mutation called in more than one member; green, mutation called in one member; gray, alleles with no call. Asterisks and colored labels mark members with a call.

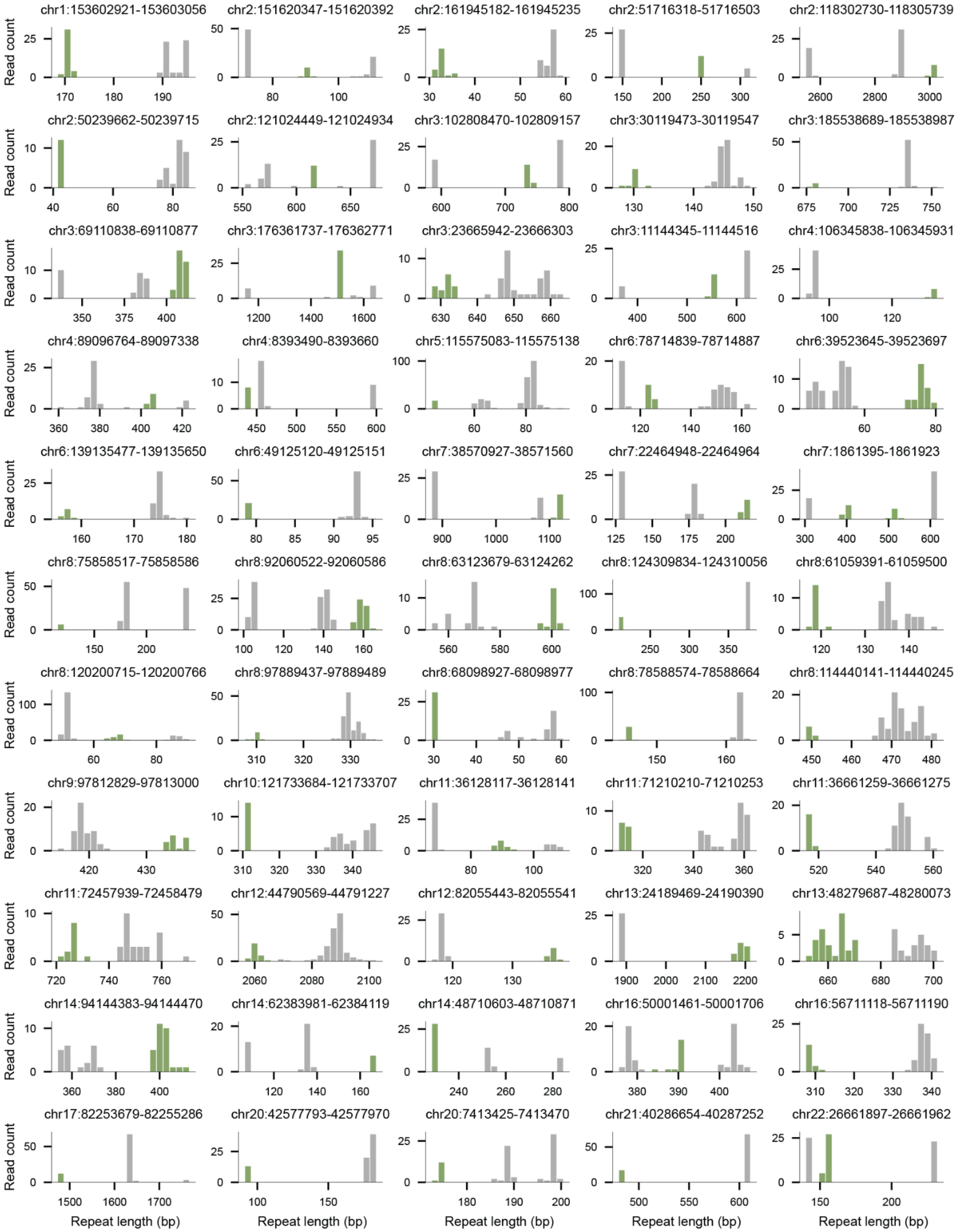

chr1:153602921-153603056
chr2:151620347-151620392
chr2:161945182-161945235
chr2:51716318-51716503
chr2:118302730-118305739
chr2:50239662-50239715
chr2:121024449-121024934
chr3:102808470-102809157
chr3:30119473-30119547
chr3:185538689-185538987
chr3:69110838-69110877
chr3:176361737-176362771
chr3:23665942-23666303
chr3:11144345-11144516
chr4:106345838-106345931
chr4:89096764-89097338
chr4:8393490-8393660
chr5:115575083-115575138
chr6:78714839-78714887
chr6:39523645-39523697
chr6:139135477-139135650
chr6:49125120-49125151
chr7:38570927-38571560
chr7:22464948-22464964
chr7:1861395-1861923
chr8:75858517-75858586
chr8:92060522-92060586
chr8:63123679-63124262
chr8:124309834-124310056
chr8:61059391-61059500
chr8:120200715-120200766
chr8:97889437-97889489
chr8:68098927-68098977
chr8:78588574-78588664
chr8:114440141-114440245
chr9:97812829-97813000
chr10:121733684-121733707
chr11:36128117-36128141
chr11:71210210-71210253
chr11:36661259-36661275
chr11:72457939-72458479
chr12:44790569-44791227
chr12:82055443-82055541
chr13:24189469-24190390
chr13:48279687-48280073
chr14:94144383-94144470
chr14:62383981-62384119
chr14:48710603-48710871
chr16:50001461-50001706
chr16:56711118-56711190
chr17:82253679-82255286
chr20:42577793-42577970
chr20:7413425-7413470
chr21:40286654-40287252
chr22:26661897-26661962
Read count
Repeat length (bp)

**Figure S14. Subclonal tandem repeat mutations in HCC1954.**

Read-level allele length distributions at loci with a subclonal call in HCC1954. Each panel gives the number of tumor reads carrying each allele length at one locus, with the subclonal allele in green and all other alleles in gray. Panel titles give locus coordinates, and lengths are in base pairs. Calls were made against the matched normal using the cancer module (Methods).

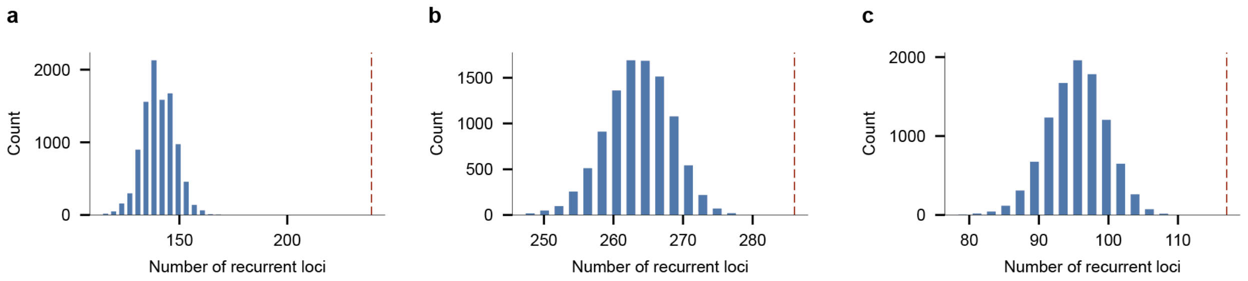


**Figure S15. Recurrence of mutated loci compared with a permutation null.**

Number of recurrent loci, those mutated in at least two samples, expected by chance. For each set of samples, the loci mutated in at least one sample were pooled, and in each of 10,000 permutations every sample was assigned a random subset of the pooled loci equal in size to its observed number of mutated loci. Histograms give the number of recurrent loci across permutations, and the dashed line marks the observed number.

a. Somatic events in six cancer cell lines, each called against its matched normal.
b. Large length changes in the 16 CEPH pedigree members in the third and fourth generation.
c. Large length changes in the same six cell lines.

**a**

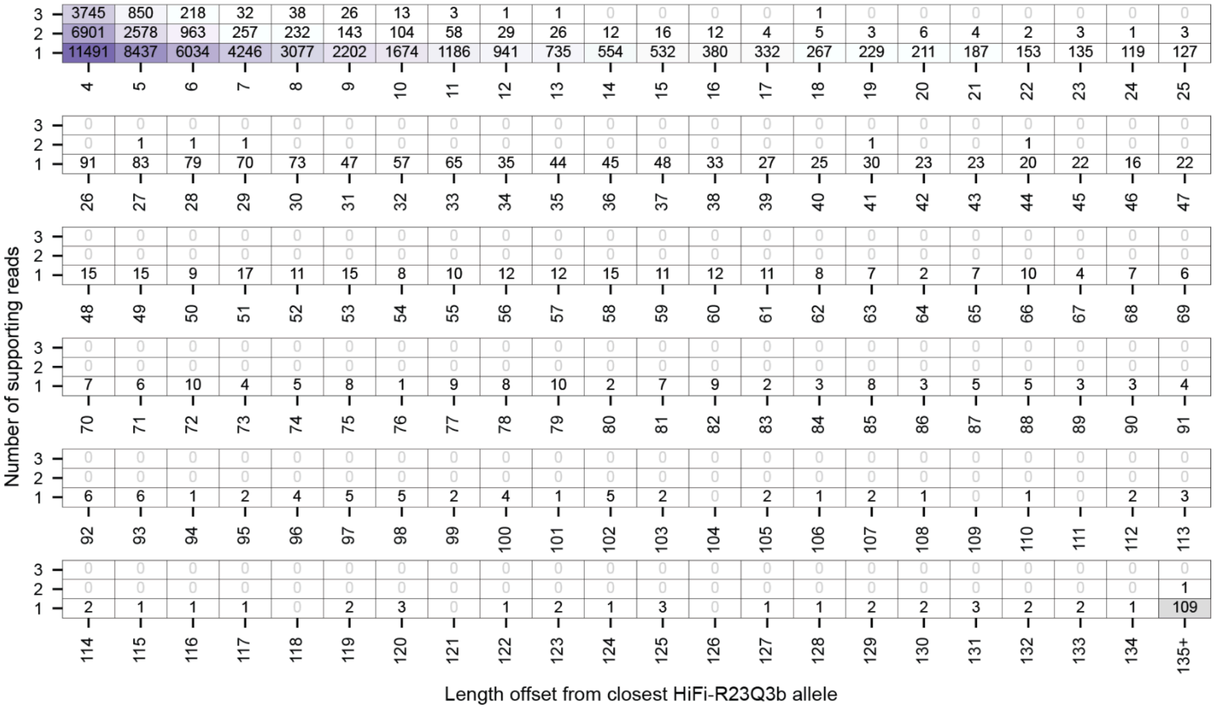
Number of supporting reads
3 | 3745 850 218 32 38 26 13 3 1 1 0 0 0 0 1 0 0 0 0 0 0 0
2 | 6901 2578 963 257 232 143 104 58 29 26 12 16 12 4 5 3 6 4 2 3 1 3
1 | 11491 8437 6034 4246 3077 2202 1674 1186 941 735 554 532 380 332 267 229 211 187 153 135 119 127
4 5 6 7 8 9 10 11 12 13 14 15 16 17 18 19 20 21 22 23 24 25
3 | 0 0 0 0 0 0 0 0 0 0 0 0 0 0 0 0 0 0 0 0 0 0
2 | 0 1 1 1 0 0 0 0 0 0 0 0 0 0 0 1 0 0 1 0 0 0
1 | 91 83 79 70 73 47 57 65 35 44 45 48 33 27 25 30 23 23 20 22 16 22
26 27 28 29 30 31 32 33 34 35 36 37 38 39 40 41 42 43 44 45 46 47
3 | 0 0 0 0 0 0 0 0 0 0 0 0 0 0 0 0 0 0 0 0 0 0
2 | 0 0 0 0 0 0 0 0 0 0 0 0 0 0 0 0 0 0 0 0 0 0
1 | 15 15 9 17 11 15 8 10 12 12 15 11 12 11 8 7 2 7 10 4 7 6
48 49 50 51 52 53 54 55 56 57 58 59 60 61 62 63 64 65 66 67 68 69
3 | 0 0 0 0 0 0 0 0 0 0 0 0 0 0 0 0 0 0 0 0 0 0
2 | 0 0 0 0 0 0 0 0 0 0 0 0 0 0 0 0 0 0 0 0 0 0
1 | 7 6 10 4 5 8 1 9 8 10 2 7 9 2 3 8 3 5 5 3 3 4
70 71 72 73 74 75 76 77 78 79 80 81 82 83 84 85 86 87 88 89 90 91
3 | 0 0 0 0 0 0 0 0 0 0 0 0 0 0 0 0 0 0 0 0 0 0
2 | 0 0 0 0 0 0 0 0 0 0 0 0 0 0 0 0 0 0 0 0 0 0
1 | 6 6 1 2 4 5 5 2 4 1 5 2 0 2 1 2 1 0 1 0 2 3
92 93 94 95 96 97 98 99 100 101 102 103 104 105 106 107 108 109 110 111 112 113
3 | 0 0 0 0 0 0 0 0 0 0 0 0 0 0 0 0 0 0 0 0 0 0
2 | 0 0 0 0 0 0 0 0 0 0 0 0 0 0 0 0 0 0 0 0 0 1
1 | 2 1 1 1 0 2 3 0 1 2 1 3 0 1 1 2 2 3 2 2 1 109
114 115 116 117 118 119 120 121 122 123 124 125 126 127 128 129 130 131 132 133 134 135+
Length offset from closest HiFi-R23Q3b allele

**b**

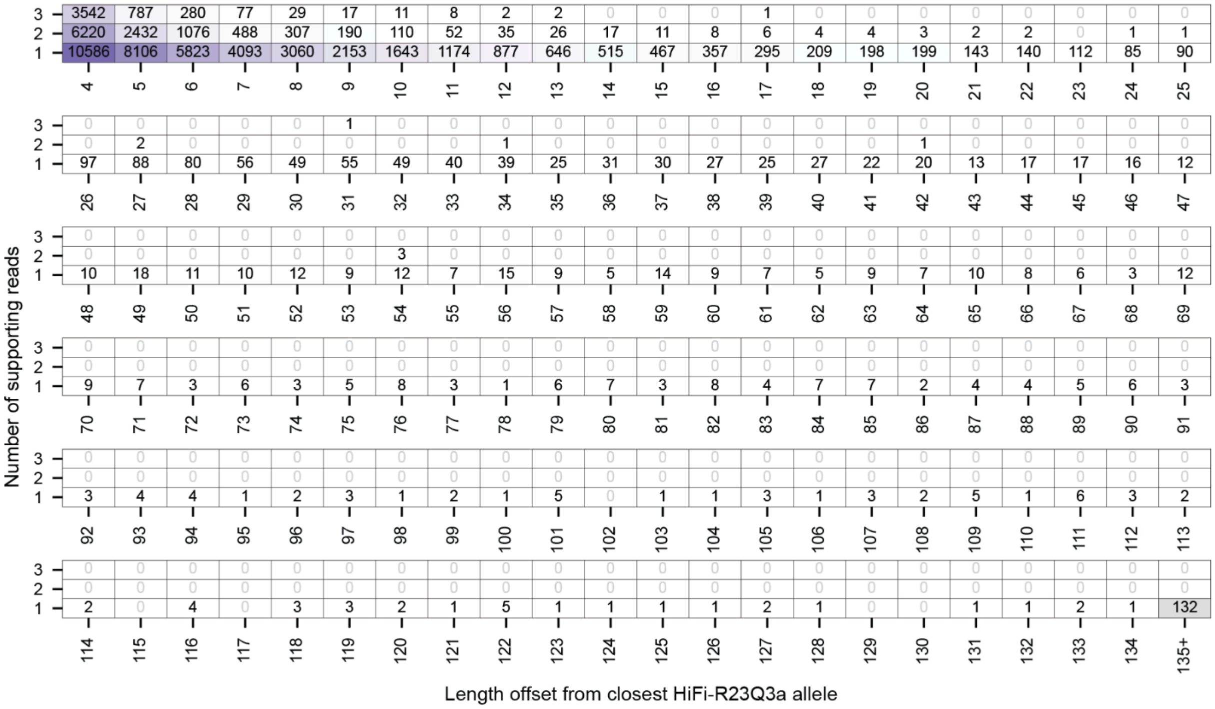
Number of supporting reads
3 | 3542 787 280 77 29 17 11 8 2 2 0 0 0 1 0 0 0 0 0 0 0 0
2 | 6220 2432 1076 488 307 190 110 52 35 26 17 11 8 6 4 4 3 2 2 0 1 1
1 | 10586 8106 5823 4093 3060 2153 1643 1174 877 646 515 467 357 295 209 198 199 143 140 112 85 90
4 5 6 7 8 9 10 11 12 13 14 15 16 17 18 19 20 21 22 23 24 25
3 | 0 0 0 0 0 1 0 0 0 0 0 0 0 0 0 0 0 0 0 0 0 0
2 | 0 2 0 0 0 0 0 0 1 0 0 0 0 0 0 0 1 0 0 0 0 0
1 | 97 88 80 56 49 55 49 40 39 25 31 30 27 25 27 22 20 13 17 17 16 12
26 27 28 29 30 31 32 33 34 35 36 37 38 39 40 41 42 43 44 45 46 47
3 | 0 0 0 0 0 0 0 0 0 0 0 0 0 0 0 0 0 0 0 0 0 0
2 | 0 0 0 0 0 0 3 0 0 0 0 0 0 0 0 0 0 0 0 0 0 0
1 | 10 18 11 10 12 9 12 7 15 9 5 14 9 7 5 9 7 10 8 6 3 12
48 49 50 51 52 53 54 55 56 57 58 59 60 61 62 63 64 65 66 67 68 69
3 | 0 0 0 0 0 0 0 0 0 0 0 0 0 0 0 0 0 0 0 0 0 0
2 | 0 0 0 0 0 0 0 0 0 0 0 0 0 0 0 0 0 0 0 0 0 0
1 | 9 7 3 6 3 5 8 3 1 6 7 3 8 4 7 7 2 4 4 5 6 3
70 71 72 73 74 75 76 77 78 79 80 81 82 83 84 85 86 87 88 89 90 91
3 | 0 0 0 0 0 0 0 0 0 0 0 0 0 0 0 0 0 0 0 0 0 0
2 | 0 0 0 0 0 0 0 0 0 0 0 0 0 0 0 0 0 0 0 0 0 0
1 | 3 4 4 1 2 3 1 2 1 5 0 1 1 3 1 3 2 5 1 6 3 2
92 93 94 95 96 97 98 99 100 101 102 103 104 105 106 107 108 109 110 111 112 113
3 | 0 0 0 0 0 0 0 0 0 0 0 0 0 0 0 0 0 0 0 0 0 0
2 | 0 0 0 0 0 0 0 0 0 0 0 0 0 0 0 0 0 0 0 0 0 0
1 | 2 0 4 0 3 3 2 1 5 1 1 1 1 2 1 0 0 1 1 2 1 132
114 115 116 117 118 119 120 121 122 123 124 125 126 127 128 129 130 131 132 133 134 135+
Length offset from closest HiFi-R23Q3a allele

**Figure S16. Replicate-unique alleles across the full range of length offsets.**

Alleles present in one HiFi replicate but absent from the other, cross-tabulated by the number of supporting reads and the length offset from the closest allele in the other replicate. Only alleles with three or fewer supporting reads are shown. Cell values and shading give allele counts, and the final column aggregates offsets of 135 bp or more.

a. Present in HiFi-R23Q3a, absent from HiFi-R23Q3b.
b. Present in HiFi-R23Q3b, absent from HiFi-R23Q3a.

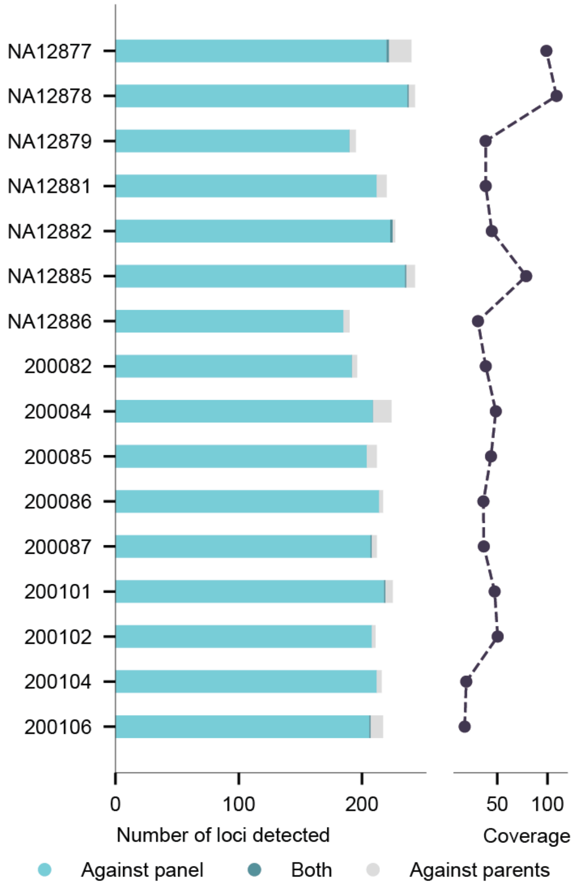


**Figure S17. Panel-based and trio-based calls in each pedigree member.**

Left, number of loci per sequenced pedigree member with an extreme allele called against the panel of normals, a mutant allele called against the member's parents with the trio module, or both. Right, sequencing coverage of each member. Only loci with an allele called in at least 10 reads in every dataset compared were analyzed, and homopolymers were excluded (Methods).

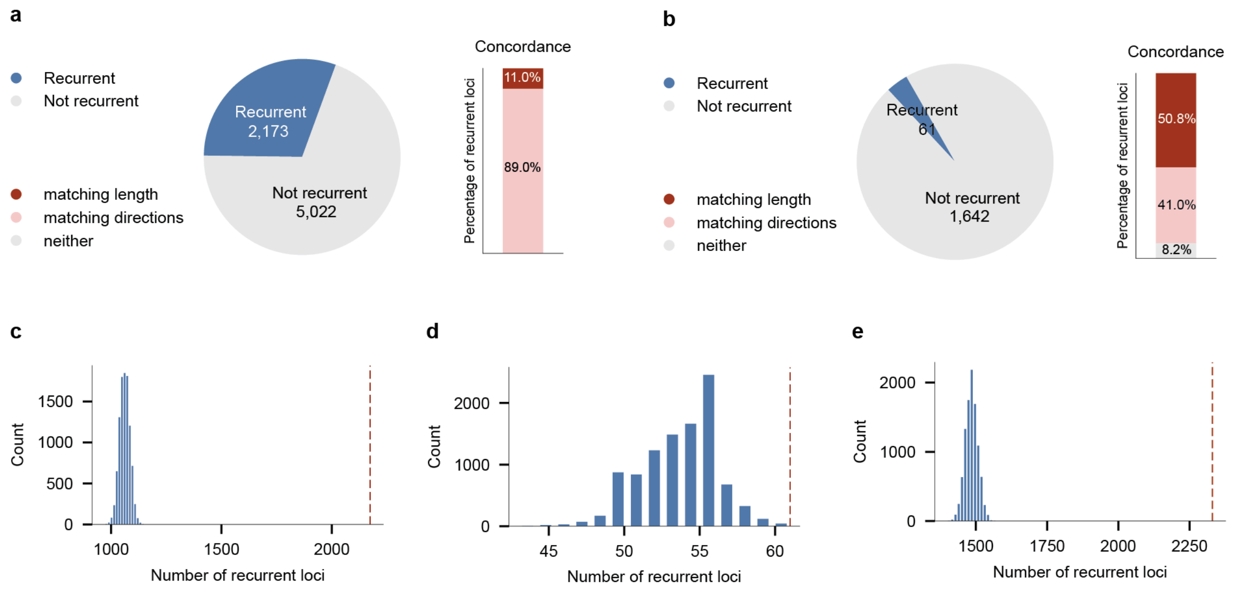


**Figure S18. Recurrence of panel-based calls across 10 cancer cell lines, by microsatellite instability status.**

Loci with an extreme allele called against the panel of normals in 10 cancer cell lines, split into two microsatellite-unstable (MSI) lines and eight microsatellite-stable (MSS) lines. Recurrent loci are those with an extreme allele in at least two lines of the set.

a. MSI lines. Left, number of recurrent and non-recurrent loci. Right, concordance of extreme alleles at recurrent loci, defined as in Fig. 6d.
b. As in a, for MSS lines.

(c-e) Number of recurrent loci expected by chance, from the permutation test described in Fig. S14, for the MSI lines (c), the MSS lines (d), and all 10 lines together (e). The histograms show the number of recurrent loci across 10,000 permutations, and the dashed line marks the observed number.

Only loci with an allele called in at least 10 reads in every dataset compared were analyzed, and homopolymers were excluded (Methods).
